# Transplantation of bacteriophages from ulcerative colitis patients shifts the gut bacteriome and exacerbates severity of DSS-colitis

**DOI:** 10.1101/2021.09.10.459444

**Authors:** Anshul Sinha, Yue Li, Mohammadali Khan Mirzaei, Michael Shamash, Rana Samadfam, Irah L. King, Corinne F. Maurice

**Author notes:** Authors contributed equally (Anshul Sinha, Yue Li).

## Abstract

Inflammatory bowel diseases (IBDs) including Crohn’s disease (CD) and ulcerative colitis (UC) are characterized by chronic and debilitating gut inflammation. Altered bacterial communities of the intestine are strongly associated with IBD initiation and progression. The gut virome, which is primarily composed of bacterial viruses (bacteriophages, phages) is thought to be an important factor regulating and shaping microbial communities in the gut. While alterations in the gut virome have been observed in IBD patients, the contribution of these viruses to alterations in the bacterial community and heightened inflammatory responses associated with IBD patients remains largely unknown. Here, we performed *in vivo* microbial cross-infection experiments to follow the effects of fecal virus-like particles (VLPs) isolated from UC patients and healthy controls on bacterial diversity and severity of experimental colitis in human microbiota-associated (HMA) mice. Shotgun metagenomics confirmed that several phages were transferred to HMA mice, resulting in treatment-specific alterations in the gut virome. VLPs from healthy and UC patients also shifted gut bacterial diversity of these mice, an effect that was amplified during experimental colitis. VLPs isolated from UC patients specifically altered the relative abundance of several bacterial taxa previously implicated in IBD progression. Additionally, UC VLP administration heightened colitis severity in HMA mice, as indicated by shortened colon length and increased pro-inflammatory cytokine production. Importantly, this effect was dependent on intact VLPs. Our findings build on recent literature indicating that phages are dynamic regulators of bacterial communities in the gut and implicate the intestinal virome in modulating intestinal inflammation and disease.

## BACKGROUND

The human gut microbiota is a complex community of microorganisms including bacteria, viruses, archaea, and various eukarya, all of which provide protection against pathogens and maintain metabolic and immunological homeostasis ^1, 2^. Intestinal bacterial communities are particularly important in guiding the appropriate development of different immune cell types and regulating the balance between pro- and anti-inflammatory responses in the gut ^2–6^. For these reasons, alterations in the gut bacteriome have been associated with various immunological disorders, including inflammatory bowel diseases (IBDs) ^7^. IBDs, comprised of Crohn’s disease (CD) and ulcerative colitis (UC), are chronic conditions in which regions of the gut are inflamed and ulcerated, often leading to debilitating abdominal pain, rectal bleeding and diarrhea. In IBD patients there is often a reduction in bacterial diversity, including a decrease in the proportion of immunoregulatory short-chain fatty acid (SCFA)-producing *Clostridia* and an increase in tissue-invasive *Enterobacteriaceae* ^7–10^. Consistent with these clinical results, several of these bacterial taxa have been shown to influence colitis severity in mouse models of intestinal inflammation ^11–15^. In addition to these changes in the gut microbiota, population-based genetic studies have revealed that several IBD risk-alleles are involved in host-microbe interactions ^16, 17^. Together, these observations have led to the general assumption that IBD is the result of an inappropriate intestinal immune response towards the gut microbiota in a genetically susceptible host. Despite our understanding of genetics and changes to the gut microbiota in IBD, the precise factors that drive bacterial alterations in IBD are not well understood.

Bacteriophages (phages), which are viruses that infect bacteria, are present at similar abundances as their bacterial hosts in the human gut and have shown to be strong regulators of bacterial communities in the mammalian gut ^18–22^. Recent work has shown that *in vivo* administration of phages in mice can alter bacterial diversity ^21–24^, disrupt bacterial interaction networks ^19^, and alter the concentration of bacterial-derived metabolites ^19^. Importantly, there is emerging data to support the idea that phages can alter disease outcomes by regulating the composition and diversity of their bacterial hosts in the gut ^21, 24, 25^.

Given the limitations of culturing gut bacteria and their associated phages, virome characterization has primarily relied on metagenomic sequencing of fecal or gut mucosal samples. However, due to the extensive diversity of phages and their low representation in databases, it has been challenging to link phages to their bacterial hosts or gain taxonomic information from viral sequence data alone ^26^ . Still, recent improvements in gut virome databases ^27, 28^ and bioinformatic tools for detecting viruses ^29^ have revealed some consistent characteristics of human gut viromes. Specifically, phage communities from healthy adults are unique ^30^, stable over time ^30, 31^, and dominated by dsDNA *Caudovirales* phages and ssDNA *Microviridae* phages ^30, 32–34^. Compared to other ecosystems, there also tends to be low virus-to-bacteria ratios (VBRs) ^20^, a high proportion of bacteria containing predicted prophages ^35^, and a high prevalence of ubiquitous crAss-like phages shown to infect *Bacteroides* in gut virome samples ^28, 30, 36^.

Accumulating evidence from metagenomic sequencing of fecal and mucosal samples indicates that the gut virome is altered in IBD patients, whether in CD or UC cohorts, and adults or children ^37–40^. Several of these reports have shown increases in the abundance and richness of the order *Caudovirales* in IBD patients ^37–39^. Some also report an increase in the relative abundance of phages predicted to infect Firmicutes, which are typically reduced in IBD and contain several species that induce anti-inflammatory immune responses ^37, 41^. Most recently, Cloone*y* et. al ^37^ showed that, in addition to shifts in viral diversity, there was also increased relative abundance of phages classified as temperate in UC and CD patients. These observations suggest that inflammatory events may initiate prophage induction in gut bacteria and a switch from the lysogenic replication cycle to lytic replication ^37^. Similar observations in mouse models of colitis further support a link between intestinal inflammation and alterations of the gut virome ^42^.

These data collectively support a relationship between IBD and intestinal phage communities. However, whether phage alterations impact gut bacterial communities, intestinal immune responses, and/or disease progression is unknown. Here, we investigated the effects of administering fecal virus-like particles (VLPs) from UC patients (UC VLPs) and non-IBD controls (healthy VLPs) to human microbiota-associated (HMA) mice, and their subsequent impact on the gut microbiome and dextran sodium sulfate (DSS)-induced colitis.

## RESULTS

### Experimental model and composition of pooled viral and bacterial stocks

In order to determine the effects of healthy and UC VLPs on bacterial and DSS-colitis severity, we performed *in vivo* “cross-infection” experiments in HMA mice. Germ-free (GF) mice were first colonized with pooled bacterial communities from 3 healthy volunteers or 3 UC patients (healthy-HMA mice, UC-HMA mice). Following bacterial colonization, mice were given single or multiple doses of VLPs, followed by 2% DSS (see experimental schematics in Fig. 1). The use of 2% DSS to induce colitis allowed for the temporal control of mild inflammation, thus enabling us to study VLP-mediated effects on bacterial communities both independent of, and in the presence of, intestinal inflammation. In addition, as DSS-induced inflammation is largely restricted to the colon, our model mimics pathology similar to that observed in UC patients ^43^.

**Figure 1.**
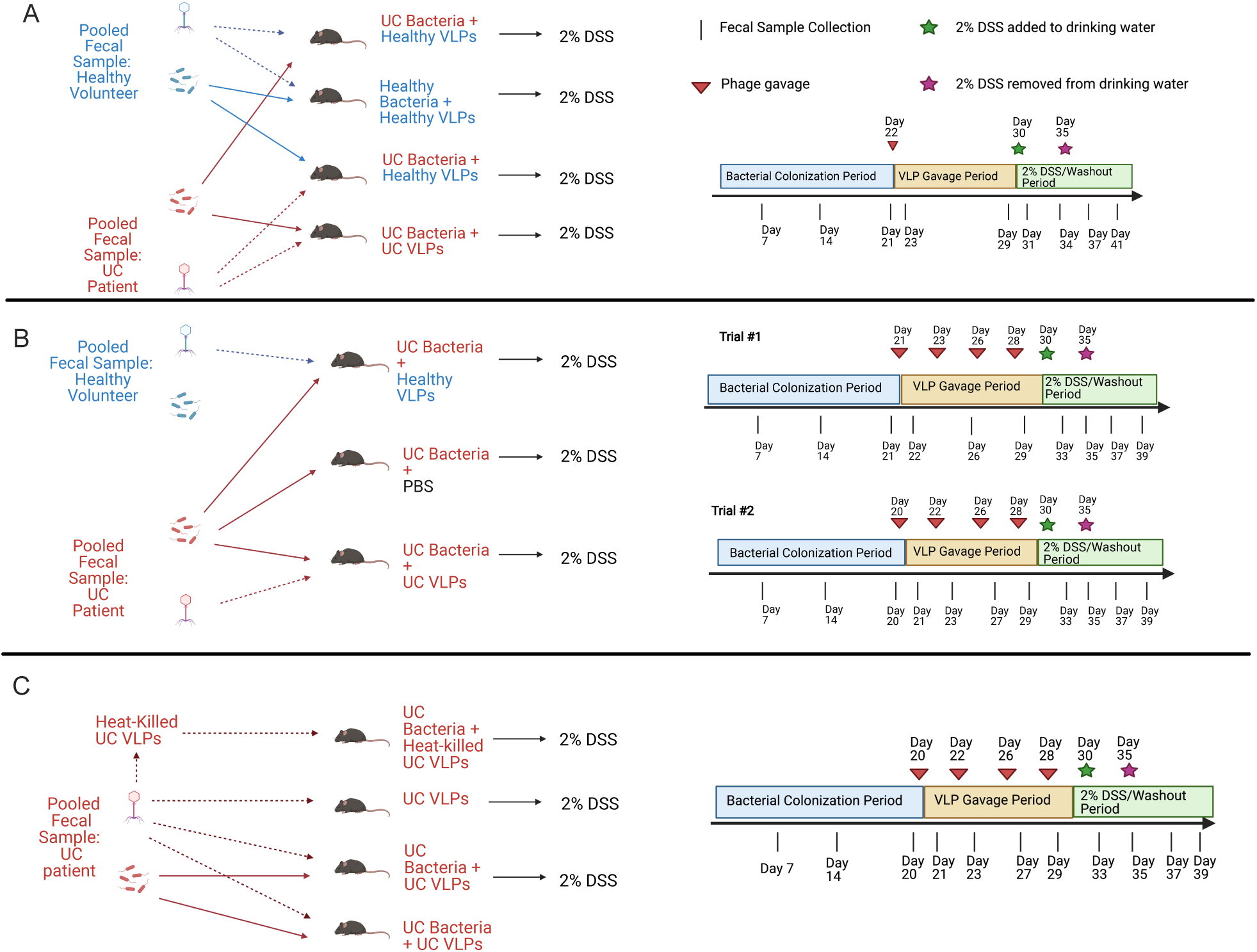
Schematic and timeline of experimental model. (A) HMA mice administered healthy or UC bacterial communities were given a single dose of healthy or UC VLPs, followed by 2% DSS. (B) UC-HMA mice were given four doses of healthy VLPs, UC VLPs or PBS, followed by 2% DSS. (C) UC-HMA or GF mice were given four doses of UC VLPs or heat-killed UC VLPs. All groups were then given 2% DSS, except one group of UC-HMA mice given UC VLPs. Each treatment group included 5 or 6 GF or HMA mice per experiment housed in three separate cages. In each experiment 200 μL of bacterial and VLP communities were administered to mice by oral gavage at equal concentrations (1-3 x 10^8^ VLPs or bacterial cells/mL).

To first characterize the virome of the pooled healthy and UC VLP stocks given to HMA mice, we performed shotgun sequencing on VLP fractions from fecal samples. Quality-filtered reads from each sample were assembled into scaffolds, and viral scaffolds were detected and annotated (see methods, Supplementary Fig. S1A). In total, 679 and 974 viral scaffolds were found in the pooled healthy and UC VLP stocks, respectively (Fig. 2A, Supplementary Table. S1). In agreement with the reported high inter-individuality of virome samples and differences in virome composition between disease states ^30, 37^, only 37 scaffolds were shared between these stock samples (Fig. 2A). We also used vConTACT2 to form viral clusters (VCs) based on shared protein-coding genetic content in order to account for the high-inter individuality of these viromes ^44^. Using this approach, 478 and 641 VCs (including singletons) were found in the healthy VLP and UC VLP stocks, respectively (Fig. 2A, Supplementary Table. S1). Of these VCs, only 89 were shared between the 2 pooled stocks (Fig. 2A), suggesting that a substantial portion of these viral stocks remained unique at this high taxonomic level. Using a vote-based approach to assign viral taxonomy to the VLP scaffolds ^30^, we also observed differences in the viral families present in each stock (Fig. 2B). Using relative abundance, the healthy stock virome was predominately composed of dsDNA phages belonging to the order *Caudovirales* and the families *Myoviridae* and *Siphoviridae* and unclassified viral scaffolds (Fig. 2B). In contrast, the pooled UC stock virome was dominated (76.36% relative abundance) by a single 6,339 bp scaffold belonging to the (ssDNA) *Microviridae* family, along with phages belonging to the *Siphoviridae* and *Podoviridae* families (Fig. 2B). These data are consistent with previous studies, showing that individual gut viromes can be dominated by ssDNA *Microviridae* ^30, 32, 34, 37^. Based on the presence of integrase, we were also able to classify viral scaffolds in our dataset as temperate ^29^. The pooled UC stock contained both higher absolute numbers of unique scaffolds identified as temperate and a higher proportion of temperate scaffolds (Fig. 2C, Supplementary Table. S1), in line with previous associations between temperate phages and IBD ^37^.

**Figure 2.**
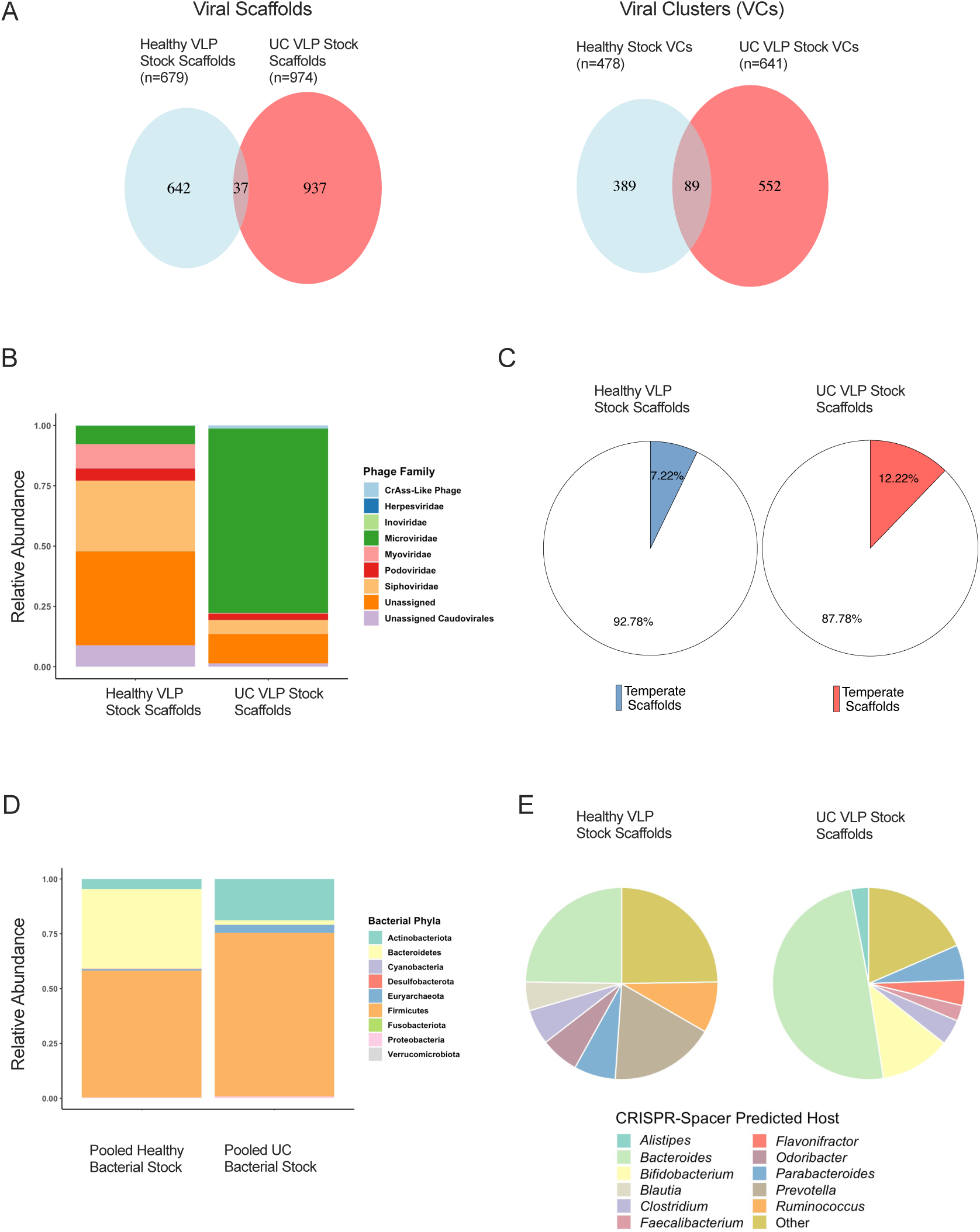
Composition of pooled healthy and UC VLP and bacterial stocks. (A) Shared and unique viral scaffolds and VCs between pooled healthy and UC VLP stocks. (B) Relative abundance of viral families in each VLP stock. (C) Proportion of scaffolds in VLP stocks identified as temperate. (D) Relative abundance of bacterial phyla in each pooled bacterial stock. (E) Proportion of scaffolds based on CRISPR spacer predicted hosts. The top 7 most prevalent host predictions in each treatment group are displayed. VLP shotgun metagenomics was used for VLP stock analyses and 16S rRNA gene sequencing of the V4 region was used for bacterial stock analyses. Stock samples were pooled using equal weight of 3 fecal samples from healthy volunteers or UC patients.

In addition to these differences in virome composition, we also used 16S rRNA gene sequencing of the V4 region to determine the composition of the pooled healthy and UC bacterial stocks used to colonize GF mice. Phylum-level analysis revealed decreased relative abundance of Bacteroidetes (2.13% UC, 36.32% healthy), increased Actinobacteria (18.80% UC, 4.63% healthy) and increased Proteobacteria (0.80 % UC, 0.15% healthy) in the pooled UC stock (Fig. 2D, Supplementary Table. S1), consistent with previous observations of UC bacterial communities ^45–47^. We also compared genus-level differences between the bacterial stocks with differences in genus-level bacterial host predictions of VLP scaffolds in our dataset using clustered regularly interspaced short palindromic repeats (CRISPR) spacer homology ^48^. In total, 186/679 (27.39 %) of healthy VLP stock scaffolds were successfully assigned CRISPR spacer-based genus predictions and 303/974 (31.11%) of UC VLP stock scaffolds were assigned genus-level predictions (Supplementary Table. S1). Some genus-level differences in relative abundance in the pooled bacterial stock were consistent with differences in genus-level bacterial host predictions of VLP scaffolds, indicative of concordance between the viral and bacterial fractions of these stocks. For instance, increased relative abundance of *Bifidobacterium* in the UC bacterial stocks (9.87% UC, 1.38% healthy) was consistent with a higher percentage of VLP scaffolds predicted to infect *Bifidobacterium* (11.88% UC, 3.23% healthy) (Fig. 2E, Supplementary Table. S1). Additionally, *Prevotella* was present at high relative abundances (21.10%) in the healthy bacterial stock and was not detected in the UC bacterial stock (Supplementary Table. S1). This disparity in *Prevotella* abundance was reflected in a high proportion of *Prevotella*-infecting VLPs in the healthy stock (17.74%) and zero scaffolds in the UC VLP stock predicted to infect *Prevotella* (Fig. 2E, Supplementary Table. S1). Interestingly, *Bacteroides*-infecting VLPs made up 49.50% of scaffolds with predicted CRISPR spacer hosts (Fig. 2E, Supplementary Table. S1), despite low *Bacteroides* relative abundance (1.79%) in the UC bacterial inoculum (Supplementary Table. S1), which could be reflective of an expansion of phages targeting and depleting *Bacteroides* in these UC patients. Together, our data highlight UC-specific alterations in both the viral and bacterial fractions of the pooled fecal samples used for VLP cross-infection experiments in HMA mice.

### UC bacterial communities enhance DSS-colitis severity in comparison to bacterial communities from healthy controls

To first determine the effects of single healthy and UC VLPs doses on bacterial diversity and DSS-colitis severity, we colonized GF mice with the pooled healthy and UC bacterial communities described above (Fig. 2D). After 21 days of bacterial colonization, mice were given a single dose of either healthy or UC VLPs, followed by 2% DSS on day 30 (Fig. 1A). After a single dose of VLPs, we did not observe differences in bacterial beta-diversity by weighted UniFrac distance between mice given healthy or UC VLPs in either healthy or UC-HMA mice (Supplementary Fig. S2A, Supplementary Table. S2). Using ANCOM, a statistical framework that accounts for the underlying structure of microbial communities ^49^, we were also unable to identify any species that were differentially abundant between mice given healthy and UC VLPs in healthy or UC-HMA mice during the VLP gavage period. Similarly, there were no significant differences in DSS-colitis severity in HMA mice given healthy or UC VLPs (Supplementary Fig. S3), suggesting that single doses of VLPs did not alter bacterial community composition or regulate intestinal inflammation.

However, regardless of whether healthy or UC VLPs were administered, UC-HMA mice had increased colitis severity compared to healthy-HMA mice as determined by innate immune cellular infiltration, inflammatory cytokine secretion in colonic explants, and tissue histology (Fig. 3A-F). Overall, these data are consistent with previous studies ^50, 51^ indicating that gut bacteria from UC patients predisposes HMA mice to an enhanced form of colitis. In order to determine the differences between the gut bacterial communities in mice humanized with microbial communities from UC patients or healthy volunteers, we performed 16S rRNA gene sequencing on mouse fecal pellets. Principal coordinate analysis (PCoA) on weighted UniFrac distances including all time points revealed significant differences in bacterial beta-diversity between healthy and UC-HMA mice (Fig. 3G, PERMANOVA p= 0.001). We next tested for treatment-specific differences in bacterial species previously associated with human IBD or experimental colitis severity. Using ANCOM, we identified 61/189 species that were differentially abundant between mice given bacterial communities from healthy volunteers and UC patients. HMA mice humanized with UC bacteria showed reduced proportions of *Akkermansia* sp. across all time points (Fig. 3H), a bacterial genus typically reduced in IBD patients and shown to ameliorate DSS-colitis ^12, 52^. It is also well established that bacteria from the *Enterobacteriaceae* family increase in abundance in IBD and exacerbate experimental colitis severity ^13, 14^. Accordingly, we found an expansion of *Escherichia*-*Shigella* sp. during DSS-colitis, only in mice humanized with UC bacteria (Fig. 3I). Importantly, in the bacterial stocks used to gavage the HMA mice, there were similar increases in *Escherichia*-*Shigella* sp. (0.45% UC, 0.069% healthy) and decreases in *Akkermansia* sp. (0 % UC, % healthy) (Supplementary Table. S1) in the UC samples compared to healthy volunteers. Together, these colonization-specific differences in bacterial taxa may explain the exacerbation of DSS-colitis in UC-HMA mice.

**Figure 3.**
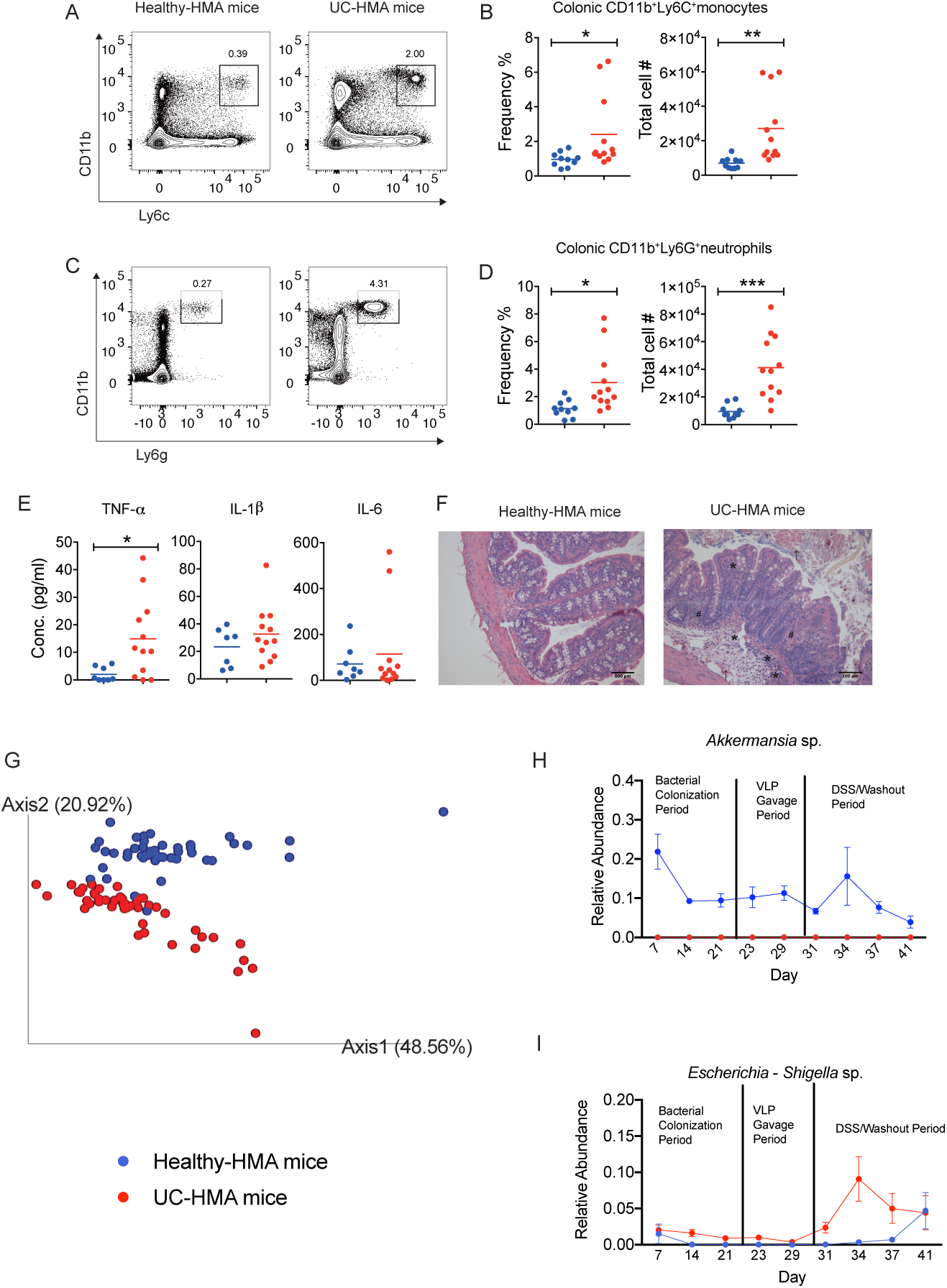
Mice colonized with UC patient-derived fecal bacteria exhibit increased inflammation during experimental colitis. (A) Representative contour plots and (B) mean frequency and absolute number of colonic inflammatory monocytes (CD11b^+^Ly6C^+^Ly6G^-^) at day 10 post-DSS administration. (C) Representative contour plots and (D) mean frequency and absolute number of colonic neutrophils (CD11b^+^Ly6C^-^Ly6G^+^) at day 10 post-DSS administration. (E) Mean TNF-α, IL-1β, IL-6 production from colon tissue explants at day 10 post-DSS administration. (F) Representative H&E staining of paraffin-embedded cross colon sections at day 10 post-DSS administration (scale bars are 100μm). Asterisk (*) indicates area of cellular infiltration; Number sign (#) indicates area of distorted crypt architecture; Black arrow indicates area of bleeding. Data were analyzed using a two-tailed unpaired parametric t test (*p < 0.05, **p < 0.01, ***p < 0.001). (G-I) 16S rRNA gene sequencing of HMA mouse fecal bacteria. (G) Beta-Diversity was determined on weighted UniFrac distances and significance was assessed using PERMANOVA (p= 0.001). (H-I) Mean relative abundance of *Akkermansia* sp. (H) *and Escherichia-Shigella* sp. (I) over time in HMA mice. Species were confirmed to be differentially abundant using ANCOM. Error bars, SE. (B, D, E) Error bars, SD. Data shown from one experiment. Dots in B, D and E represent individual mice (n=8 healthy-HMA mice, n=12 UC-HMA mice). Dots in G, H, I indicate pooled mouse fecal samples at a single time point. At each time point, mouse fecal samples in each cage were pooled from 1 or 2 mice (n=6 cages per group, 1 or 2 mice per cage).

### Administration of healthy and UC VLPs increases fecal viral abundance and virus-to-bacteria ratio (VBR) in HMA mice

We next wanted to determine whether multiple VLP doses from healthy volunteers could alter the gut bacteriome and prevent the exacerbation of DSS-colitis. As we did not observe noticeable changes in bacterial community composition (Supplementary Fig. S2A, Supplementary Table. S2) or DSS-colitis severity (Supplementary Fig. S3) following a single dose of healthy or UC VLPs, we explored whether multiple doses of healthy or UC VLPs would alter the gut microbiota of UC- HMA mice (Fig. 1B-C). One common approach in phage therapy to increase treatment efficacy is to add multiples doses instead of one single dose to sustain high phage densities ^53^. Thus, for the remaining experiments, we proceeded to give HMA mice multiple repeated doses of VLPs.

In the first experiment (Fig. 1B), UC-HMA mice were given four doses of healthy VLPs, UC VLPs, or PBS over the course of 9-10 days. Following the VLP gavage period, mice were given 2% DSS followed by a washout period. This first experiment was performed independently twice (two trials). In a second experiment (Fig. 1C), we tested the impact of phage viability as well as bacteria-independent effects of multiple doses of VLPs on intestinal inflammation using UC- HMA mice administered intact or heat-killed UC VLPs and GF mice given UC VLPs alone (Fig. 1C). We first determined whether repeated dosing of VLPs would increase viral abundance and virus-to-bacteria ratio (VBR) in HMA mice. Low levels of VLPs (8.12 x 10^8^ virus mL^-1^) could be detected in HMA mice during the bacterial colonization period (Fig. 4A) before the addition of stock VLPs, likely a result of prophage induction of newly colonized bacteria and/or VLPs that we were unable to remove during separation of the bacterial and VLP fractions in the stocks. Still, these levels of VLPs during the bacterial colonization period were lower compared to after VLP gavage (3.93-fold increase in UC VLP treated mice and 3.91-fold increase in healthy VLP treated mice) (Fig. 4A). Additionally, regardless of the source (healthy or UC samples), repeating VLP dosing resulted in a significant increase in viral abundance and virus-to-bacteria ratio (VBR) during the VLP gavage period relative to PBS (Fig. 4A-D, Supplementary Fig. 4A-D) or heat-killed UC VLP controls (Supplementary Fig. 4E-H). Despite this, there was no associated decrease in total bacterial abundance compared to PBS (Fig. 4B, Supplementary Fig. S4B) or to heat-killed controls (Supplementary Fig. S4F), which suggests that bacteria killed by phage-mediated lysis may be rapidly replaced by genetically or phenotypically resistant bacterial taxa ^19, 54^. Notably, there was no detectable increase in viral abundance in GF mice given UC VLPs alone, suggesting that phage replication required bacterial hosts (Supplementary Fig. S4E).

**Figure 4.**
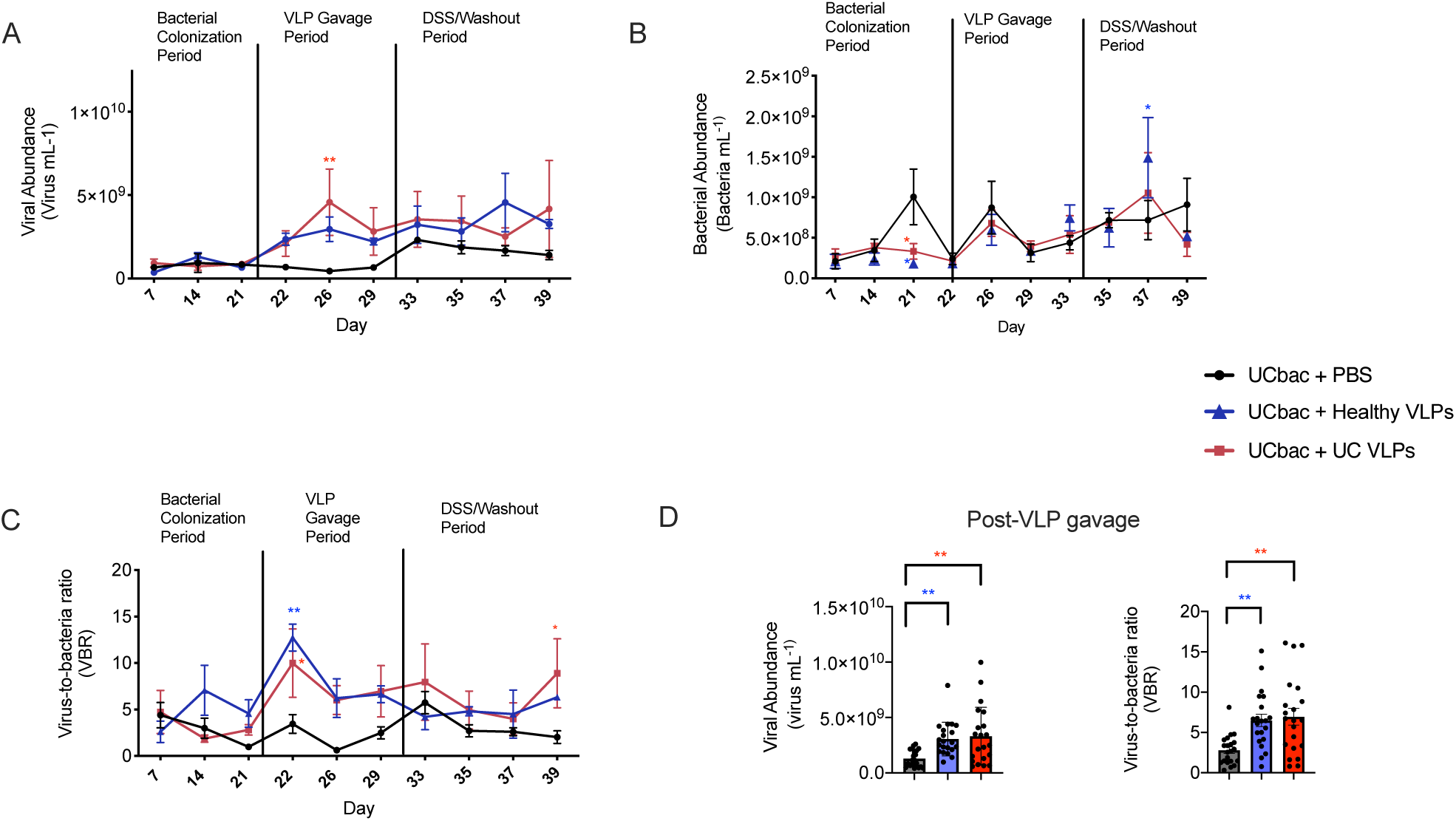
Healthy and UC VLP administration increases viral abundance and VBR in UC-HMA mice. (A) Viral abundance was determined from mouse fecal pellets using epifluorescence microscopy and compared to (B) bacterial abundances obtained by flow cytometry after staining with SybrGREEN I to obtain (C) VBRs. (D) Mean total viral abundance and mean total VBR post-VLP gavage were compared between treatment groups after the first dose of VLPs or PBS was given to mice. (A-C) Significance was assessed using two-way ANOVA and Dunnett’s multiple comparisons test (*p ≤ 0.05, **p≤ 0.01). (D) Significance was assessed using one-way ANOVA and Tukey’s multiple comparison test (**p≤ 0.01). Red and blue asterisks indicate significant differences between the PBS control and HMA mice given UC VLPs and healthy VLPs, respectively. Data shown is from one of two independent trials. Dots represent abundance or VBR of pooled mouse fecal samples at a single time point. At each time point, mouse fecal samples in each cage were pooled from 2 mice (n=3 cages per group, 6 mice per group). Error bars, SE. UC bac, UC-HMA mice.

Since some ecological models predict that more metabolically active bacteria are less susceptible to lytic phage infection ^55^, we next determined if healthy and UC VLPs could alter the proportion of active bacterial cells. Using SYBRGreen I staining and flow cytometry, we determined the proportion of active fecal bacteria in HMA mice, as described previously ^56^ (Supplementary Fig. S5A). Compared to PBS and heat-killed UC VLP controls, healthy and UC VLPs did not shift the proportion of active bacteria (Supplementary Fig. S5C, S5E, S5G). We also used Propidium Iodide (PI) staining to determine the proportion of damaged bacterial cells in HMA mice (Supplementary Fig. S5B). Healthy and UC VLPs also did not shift the proportion of damaged cells compared to heat-killed or PBS controls, with the exception of a single time-point, where UC VLPs increased the proportion of damaged bacteria (Supplementary Fig. S5D, S5F, S5H). Together, these data suggest that donor VLPs can interact with HMA mice but are not able to shift the proportions of active or damaged gut bacterial cells to a detectable level.

### Fecal VLPs derived from healthy volunteers and UC patients drive unique changes in the virome of HMA mice

Shotgun sequencing was also performed on the VLP fraction of mouse fecal pellets to follow changes in virome composition of HMA mice following VLP gavage. To assess the effectiveness of transfer of VLPs from the pooled stocks to the HMA mice, we determined the proportion of scaffolds found in the VLP stocks that were only found in HMA mice post-VLP gavage. In total, 82/679 (12.06%) of healthy VLP stock scaffolds were also found in HMA mice post healthy VLP gavage (Supplementary Table. S3). In contrast, a higher number and proportion of UC VLP stock scaffolds, reaching 210/974 (21.56%), were found in HMA mice post UC VLP gavage, possibly reflective of autologous transfer of UC VLPs to UC-HMA mice (Supplementary Table. S3). To further assess the effectiveness of VLP transfer, we performed non-metric multidimensional scaling (NMDS) on Jaccard distance of VCs and we measured the Jaccard distance of VCs between each stock and HMA samples before and after transfer to HMA mice. Compared to the viromes of mice before VLP gavage, there was a significant reduction in Jaccard distance between the viromes of pooled healthy stock and the UC-HMA mice given healthy VLPs, which suggests that phages present in the healthy stock are persisting in these mice after gavage (Fig. 5A, left). The same comparison was made using UC VLPs without significant differences observed (Fig. 5A, right). In line with these data, after VLP gavage, there was a significant increase in the richness of viral scaffolds and VCs in UC-HMA mice given healthy VLPs and a non-significant increase in richness of viral scaffolds in UC-HMA mice given UC VLPs compared to PBS controls (Supplementary Fig. S1B).

**Figure 5.**
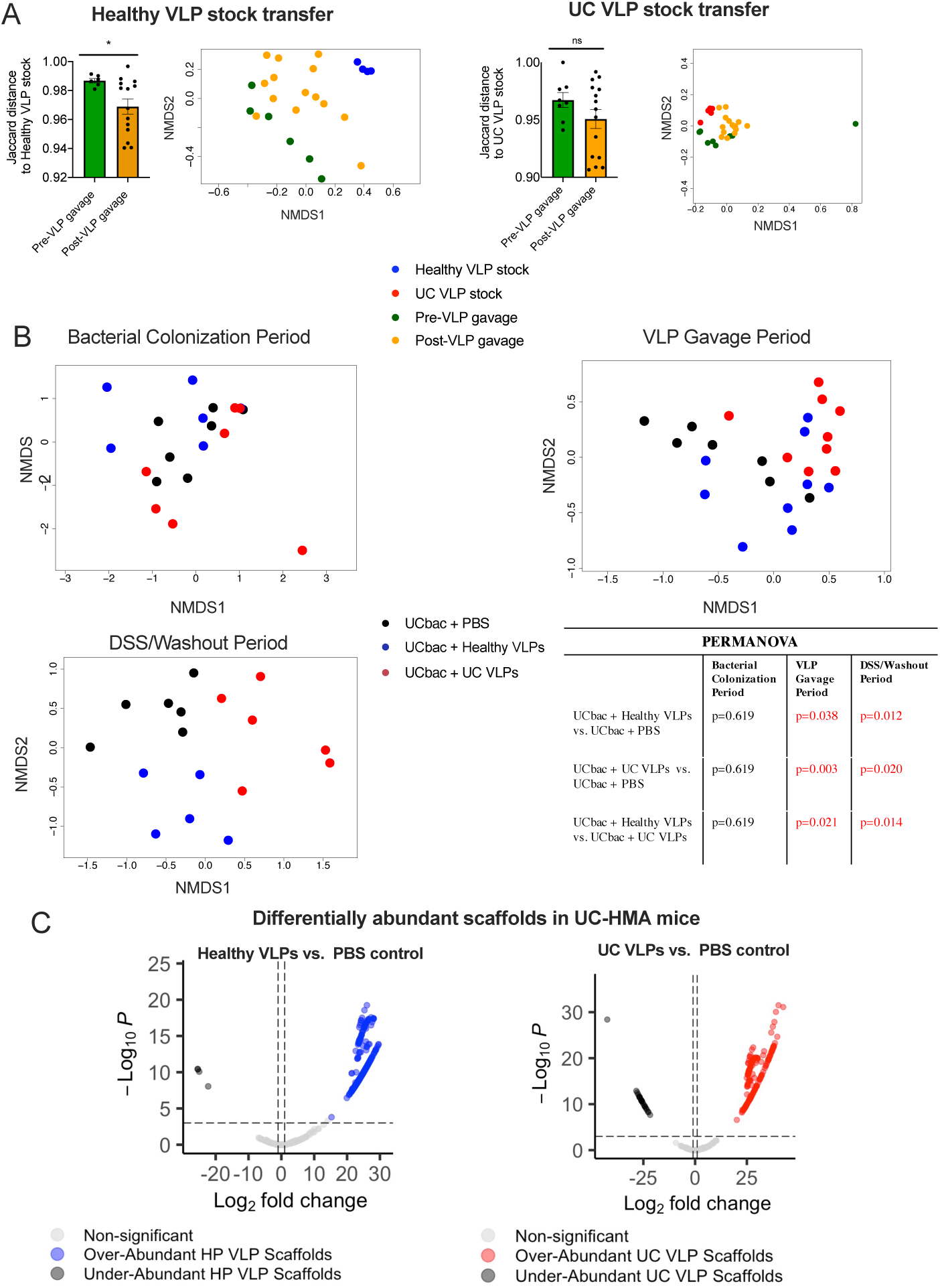
Differences in virome composition of UC-HMA mice after VLP gavage. (A) NMDS of Jaccard distance and quantification of Jaccard distance of VCs to the healthy VLP stock (left) and UC VLP stock (right) in HMA samples before and after VLP gavage. Post-VLP gavage samples included all samples after the first dose of VLPs were added. Significant differences in Jaccard distances were assessed using an two-tailed unpaired t-test (p* ≤ 0.05). Error bars, SE. (B) NMDS of Bray-Curtis dissimilarity of scaffolds between HMA mice given healthy VLPs, UC VLPs or PBS during the (top-left) bacterial colonization period, (top-right) VLP gavage period or (bottom-left) DSS washout period. Significant differences in Bray-Curtis dissimilarity were assessed in each time period using adonis PERMANOVA (p ≤ 0.05) (bottom-right). Dots represent pooled mouse fecal samples at a single time point. Samples from all time points of the longitudinal study were included in the NMDS and comparative analyses. (C) DESeq2 differentially abundant scaffolds between UC-HMA mice given healthy (left) and UC (right) VLPs. Dots shown in the volcano plots represent the differentially abundant scaffolds found during the VLP gavage period. Scaffolds with adjusted p-values < 0.001 and with log2fold changes greater or less than 1 were considered differentially abundant using a two-tailed wald test ^85^. Data shown are from the first of two independent trials represented in Fig. 1B (trial #1). At each time point, mouse fecal samples in each cage were pooled from 2 mice (n=3 cages per group, 6 mice per group. UCbac, UC-HMA mice).

We next performed NMDS on Bray-Curtis dissimilarity of viral scaffolds and VCs to follow differences in viral beta-diversity over time in UC-HMA mice. As expected, there were no significant differences in Bray-Curtis dissimilarity during the baseline bacterial colonization period (Fig. 5B, Supplementary Fig. S1C). In all cases, however, significant differences in viral scaffold diversity were observed in all groups given VLPs compared to PBS controls, further indicating successful transfer and replication of donor VLPs (Fig. 5B). Similar trends were observed with VCs, however, there were no significant differences observed between healthy VLP-treated mice and PBS controls during the VLP gavage period (Supplementary Fig. S1C, p=0.054). Notably, significant differences in viral diversity between all groups were maintained during the DSS washout period (Fig. 5B, Supplementary Fig. S1C). To further assess differences between VLP-treated mice and PBS controls, we performed differential abundance analysis of viral scaffolds using DESeq2 ^57^. During the VLP gavage period prior to DSS administration, there were 196 differentially abundant viral scaffolds between UC VLP-treated mice controls and PBS controls (173 over-abundant, 23 under-abundant) and 272 differentially abundant scaffolds between healthy VLP-treated mice and PBS controls (268 over-abundant, 4 under-abundant) (Fig. 5C, Supplementary Table. S4). During the DSS washout period, there were 303 differentially abundant scaffolds between UC VLP-treated mice (209 over-abundant, 94 under-abundant) and PBS controls and 232 differentially abundant scaffolds between healthy VLP-treated mice and PBS controls (168 over-abundant, 64 under-abundant) (Supplementary Table. S4). Of these significantly upregulated scaffolds, 46/173 (VLP gavage period) and 60/209 (DSS washout period) in UC VLP-treated mice and 23/268 (VLP gavage period) and 34/168 (DSS washout period) in healthy VLP-treated mice were also found in their respective pooled inoculums. Of the most over- abundant scaffolds with CRISPR spacer-predicted hosts, mice given healthy VLPs were enriched with *Bacteroides*-infecting phages during both the VLP gavage and DSS washout periods (Supplementary Table. S4). In contrast, UC VLP-treated mice were enriched with phages predicted to infect *Hungatella, Barnsiella*, *Erysipelatoclostridium,* and *Bacteroides* during the VLP gavage period and were enriched with *Bacteroides, Faecalibacterium* and *Eubacterium* phages during the DSS washout period (Supplementary Table. S4). Together, these data indicate that VLPs were transferred from pooled stocks to HMA mice, with distinct changes in viral beta-diversity and potential phage-bacterial interactions following VLP gavage.

### VLPs derived from healthy individuals or UC patients have distinct effects on the UC gut bacterial communities in vivo

In order to identify whether multiple healthy and UC VLPs doses had distinct effects on bacterial community composition, we performed 16S rRNA gene sequencing on mouse fecal pellets before and after VLP administration, as well as during the DSS/washout period. Using ANCOM, we identified bacterial species differentially abundant between UC-HMA mice given multiple doses of either UC VLPs, healthy VLPs, or PBS (summarized in Supplementary Tables. S5-8). To account for variation due to isolator-specific differences during colonization, bacterial species that were differentially abundant during the baseline bacterial colonization period were not included. In addition, the remaining differentially abundant species were each ranked based on the likelihood that treatment-specific differences were due to VLP treatment or isolator effect (1: likely due to VLP treatment, 2: possibly due to VLP treatment, 3: likely due to isolator effect, see methods for ranking criteria).

Across the two trials where UC-HMA mice were given multiple doses of healthy VLPs, UC VLPs, or PBS, 11 differentially abundant species (4/90 species trial #1; 7/85 species trial #2) were identified in the VLP gavage period before DSS administration, and 19 species (4/94 species trial #1; 15/85 species trial #2) were identified in the DSS/washout period (Supplementary Tables. S5-6). Between mice given multiple doses of UC VLPs (+/- DSS) or heat-killed UC VLPs, 3/49 differentially abundant bacterial species were identified in the VLP gavage period and 6/47 species were identified in the DSS/washout period (Supplementary Tables. S7-8). Some of these differentially abundant bacterial species have been shown to influence experimental colitis severity, or to be differentially abundant in CD or UC patients. For example, *Eubacterium limosum* was significantly reduced in mice given UC VLPs (Fig. 6A, left), consistent with its ability in mouse models to ameliorate DSS-colitis severity ^58^. We also observed that *Enterococcus* sp., which have been shown to play divergent roles in the context of DSS-colitis severity ^59, 60^, were increased in relative abundance in HMA mice (Fig. 6A, middle). In addition, compared to heat-killed UC VLPs, mice given intact UC VLPs showed a significant increase in the proportion of *Escherichia-Shigella* sp. after VLP gavage (Fig. 6A, right). Given that *Enterobacteriaceae* are increased in IBD patients and are thought to exacerbate dysregulated immune responses in IBD ^7, 13, 61^, these data suggest that UC VLPs may allow for the expansion of this species.

**Figure 6.**
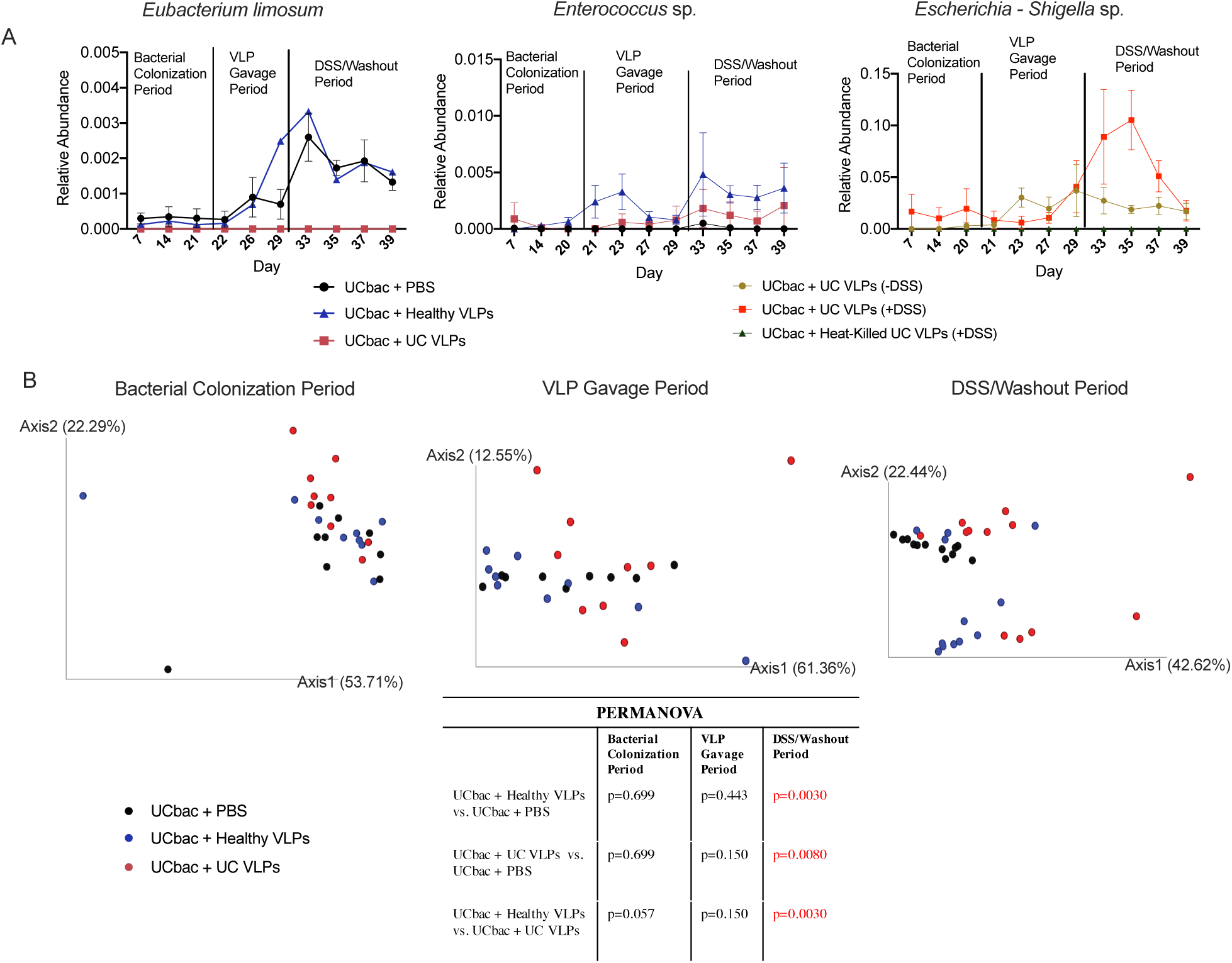
UC VLP administration differentially modulates bacterial community composition in UC-HMA mice. (A) Mean relative abundance of *Eubacterium limosum* (left), *Enterococcus* sp. (middle) and *Escherichia-Shigella* sp. (right) in UC-HMA mice over time was determined using 16S rRNA gene sequencing. ANCOM was used to confirm that these species were differentially abundant between treatment groups ^49^. Error bars, SE. (B) PcoA on weighted UniFrac distance between HMA mice during the bacterial colonization period (left), VLP gavage period (middle) and DSS/washout period (right). Significant differences between weighted UniFrac distances were assessed using pairwise PERMANOVA (p ≤ 0.05) (bottom). PcoA and weighted UniFrac distances shown are from the first of two independent trials represented in Fig. 1B (trial #1). Samples from all time points were included in the PcoA and comparative analyses. Mouse fecal samples in each cage were pooled from 2 mice (n=3 cages, 6 mice per treatment group). Dots represent pooled mouse fecal samples at a single time point. UCbac, UC-HMA mice.

PCoA on weighted UniFrac distances revealed no significant differences in bacterial beta-diversity of HMA mice during the bacterial colonization period and during the following 9-day period where 4 doses of VLPs (healthy, UC, or PBS) were given to these mice (Fig. 6B). In contrast, we observed significant differences in bacterial community composition between mice given healthy VLPs, UC VLPs, or PBS following DSS administration, supporting the idea that these phages have distinct effects on bacterial communities, which are amplified during DSS colitis (Fig. 6B). Similar trends were observed in a second independent trial (Supplementary Fig. S2B, Supplementary Table. S2). In line with these data, there were significant differences in bacterial diversity between UC-HMA mice given UC VLPs (+DSS) compared to those given heat-killed UC VLPs (+DSS) (Supplementary Fig. S2C, Supplementary Table. S2), suggesting that intact viruses are necessary for driving changes in bacterial diversity. In contrast, we did not observe differences in weighted UniFrac distances during the DSS/washout period between UC- HMA mice given UC VLPs in the absence of DSS, compared to heat-killed UC VLPs (Supplementary Fig. S2C, Supplementary Table. S2). These data further support the idea that phage-mediated changes in gut bacterial communities are initially modest, but then amplified during intestinal inflammation. In order to investigate whether these greater changes in bacterial diversity following DSS administration were in part due to prophage induction as a result of intestinal inflammation, we also determined the proportion of VLP scaffolds identified as temperate following DSS. Within each treatment group, there was a modest increase in the relative abundance of VLP scaffolds identified as temperate following DSS (Supplementary Fig. S1D). We also identified significantly over-abundant scaffolds after DSS-administration within each treatment group that were classified as temperate. Despite the modest increase in temperate VLP relative abundance following DSS, only 1/16 (UC VLP treatment), 3/8 (healthy VLP treatment) and 8/30 (PBS control) of the over-abundant scaffolds were identified as temperate (Supplementary Table. S4), suggesting that any DSS-mediated effects on prophage induction are likely modest. To determine whether there was experimental evidence for prophage induction due to direct interactions between DSS and UC bacterial communities, we performed an *in vitro* prophage induction assay ^62–64^ . In UC bacterial cultures grown anaerobically in the presence of DSS, we did not detect an increase in VLPs in the bacterial supernatant (Supplementary Fig. S1E), suggesting that any DSS-mediated prophage induction occurring in HMA mice is likely not due to direct DSS toxicity to bacterial cells and is likely through DSS-mediated induction of downstream immune responses.

Together, these data indicate that healthy and UC VLPs drive distinct changes in the relative abundance of bacterial species in UC-HMA mice, some of which have been shown to influence experimental colitis severity and IBD disease progression. In addition, while several bacterial species were found to be differentially abundant during the VLP gavage period, differences in bacterial diversity between VLP-treated mice are greatest during DSS-colitis, suggesting that intestinal inflammation may provide a more conducive environment for phage-mediated changes in these gut bacterial communities.

### UC VLPs increase colitis severity in HMA mice

Given that the healthy and UC VLPs are distinct in their composition and contribute differently to bacterial community composition, we next tested if they had distinct effects on colitis outcomes. We first determined whether there were differences in colitis severity between UC-HMA mice given healthy VLPs, UC VLPs, or PBS. Upon DSS administration, a slight decrease of body weight could be seen only in mice administered UC VLPs (Fig. 7A). Importantly, at day 10 post-DSS challenge, only the weight of UC-HMA mice given UC VLPs was declining (Fig. 7A). HMA mice given UC VLPs also showed a significant shortening of colon length (Fig. 7B) and an increase of pro-inflammatory cytokine production (Fig. 7C), indicating that UC VLPs can exacerbate the severity of DSS colitis compared to healthy VLPs and the PBS control.

**Figure 7.**
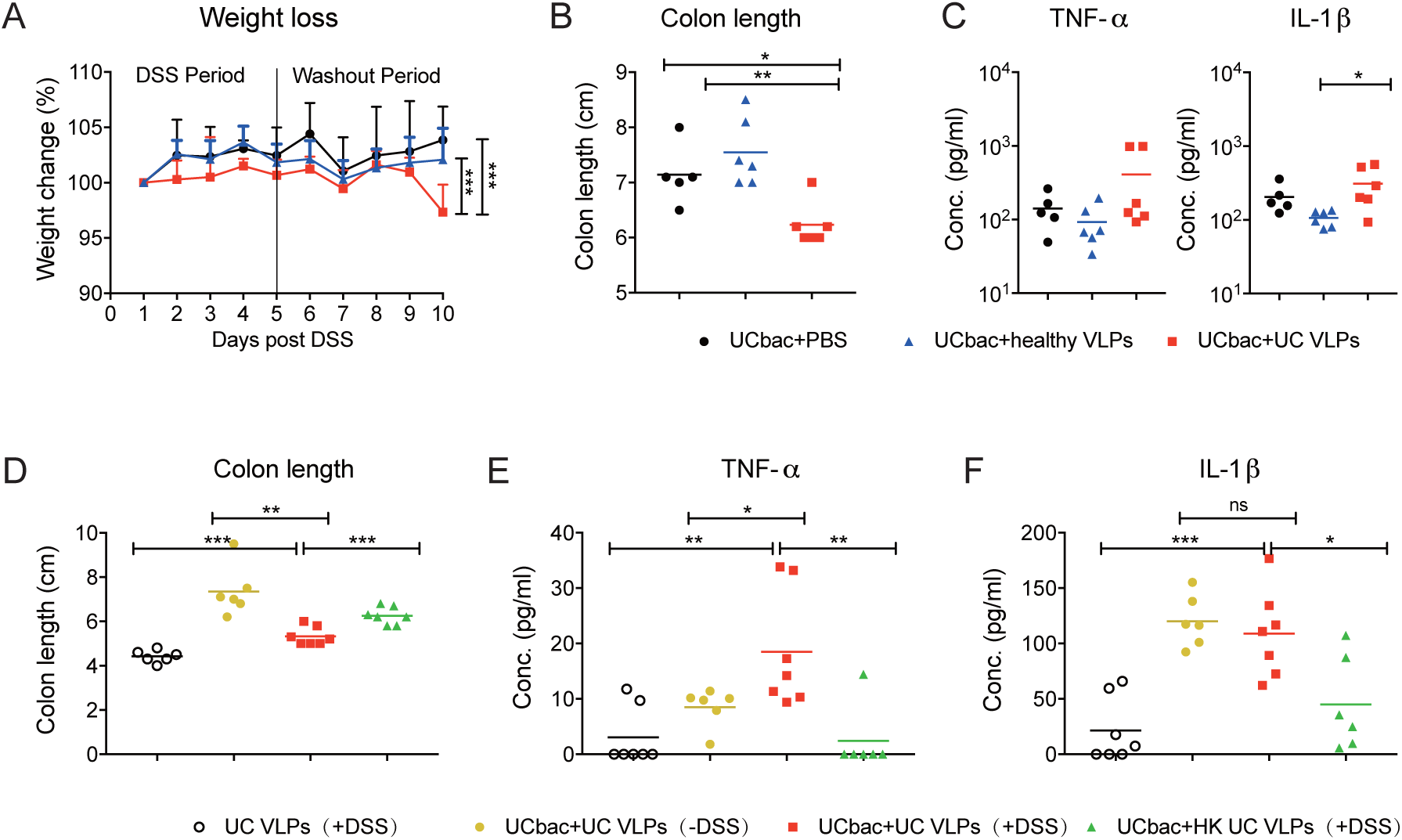
UC VLPs exacerbate the severity of experimental colitis in the presence of gut bacteria. (A) Mean body weight change during experimental colitis induction between the indicated groups. (B) Mean colon length at day 5 post-DSS administration between the three groups. (C) Mean inflammatory cytokine production in colon tissue explant at day 5 post-DSS administration. (D) Mean length of colon at day 5 post-DSS administration of the following groups: UC bacteria colonized mice treated with UC VLPs with/without DSS challenge, UC bacteria colonized mice treated with heat-killed (HK) phages and UC VLP treatment alone. (E-F) Mean inflammatory cytokine production in colon tissue explants of the indicated groups. (A) Data were analyzed by two-way ANOVA with Bonferroni for multiple comparisons. (B-F) Data were analyzed using a two-tailed unpaired parametric t test (*p < 0.05, **p < 0.01, ***p < 0.001). Error bars, SD. Data shown from one experiment. Each dot represents an individual mouse. UCbac, UC-HMA mice.

To further investigate the function of UC VLPs on the pathology of DSS-induced colitis, we next assessed DSS-colitis severity between UC-HMA mice given (i) UC VLPs with DSS; (ii) UC VLPs without DSS to determine if UC VLPs can cause spontaneous colitis; (iii) heat-killed UC VLPs to determine if intact VLPs are necessary for increased colitis severity; and (iv) GF mice given UC VLPs to determine if increased colitis severity was a result of direct UC VLP-immune interactions (Fig. 7C).

Consistent with our previous results (Fig. 7C), UC-HMA mice given UC VLPs had a significantly shortened colon and produced the most TNF-α and IL-1β from colon explants compared to the other UC-HMA groups (Fig. 7D-F). Notably, the heat-killed UC VLP group mice showed significantly less severe colitis compared to the intact UC VLPs group, suggesting the changes we observe are due to viral infection rather than the presence of phage components alone (Fig. 7D-F). Finally, we found that GF mice given UC VLPs prior to DSS treatment exhibited the greatest colon shortening, yet least inflammatory cytokine production (Fig. 7D-F). These results are similar to our observations of increased colonic shortening in GF mice compared to HMA mice (Supplementary Fig. S6) and suggest that, without bacterial hosts, phages cannot modulate colitis outcomes.

## DISCUSSION

Phages are known regulators of bacterial diversity and metabolism in several environments, including the mammalian gut ^18, 65, 66^. In IBD, metagenomic analyses have revealed that the virome composition of CD and UC patients is altered in comparison to non-IBD controls ^37–40^. However, experimental work investigating the interactions between phages, intestinal bacterial communities, and immune responses in the context of IBD is limited ^67^. Given the importance of gut bacterial communities in human health, phage-mediated changes of the microbiota are thought to have important physiological consequences ^26^. Here we show that VLPs isolated from healthy volunteers and UC patients differentially modulate the composition of the gut microbiota, and that administration of multiple doses of UC VLPs increases colitis severity in UC-HMA mice.

Four doses of UC VLPs and healthy VLPs significantly altered the fecal bacteriome of UC- HMA mice compared to PBS and heat-killed UC VLP-treated controls. This was not seen when a single dose of UC VLPs was used, supporting the idea that the regulatory effect of phages in the mammalian gut is dose dependent ^23^. The differences in the effects of multiple doses compared to single doses of VLPs on bacterial community composition and downstream disease outcomes have important implications for targeted or whole community phage therapy approaches in the gut ^26^.

Several bacterial species were found to be differentially abundant during the VLP gavage period prior to DSS-colitis, suggesting that VLPs had distinct effects on the bacterial community in the absence of induced inflammation. However, we only observed significant differences in bacterial beta-diversity during the DSS/washout period, highlighting that these initial VLP-mediated effects are amplified during the DSS-colitis/washout period. The disrupted colonic mucus layer, potential changes in bacterial growth rate, and/or innate immune cell-derived mediators in the inflamed gut could all provide a more conducive environment for increased phage- host interactions ^55^. Given these drastic changes in the gut environment during inflammation, it is possible that this facilitates a switch from Piggyback-the-Winner dynamics, common in the mammalian gut, to Kill-the-Winner dynamics ^55^. Changes in microbial expression profiles in IBD patients ^68^ and in murine experimental colitis ^69^ could also alter bacterial susceptibility to phage infection.

The use of DSS-colitis as a model of inflammation is limited in that the immune responses elicited and the exaggerated damage to the epithelium differ from clinical UC ^70^. However, we induced a mild colitis, as HMA mice given 2% DSS experienced minimal weight loss and blood in their stool. In addition, the reproducibility and temporal control offered by DSS were valuable in our approach to study the effects of VLPs on bacterial communities in the context of broad inflammatory progression and changes. Thus, our data provide the framework for future studies investigating phage-host interactions during inflammation. Given the modest increase in the relative abundance of temperate phages observed across treatment groups following DSS, prophage induction resulting from DSS-induced inflammation may also contribute to these differences in bacterial diversity. Only a small fraction of differentially abundant scaffolds were identified as temperate during the DSS washout period, yet we are likely underestimating this number since all samples collected during DSS exposure could not be used for VLP shotgun sequencing due to DSS-mediated inhibition of DNA polymerase required for multiple displacement amplification (MDA), despite our attempts to purify DNA using spermidine ^71^. Interestingly, while we did observe differences in bacterial community diversity based on VLP treatment, there were no differences in bacterial activity or damage in VLP treated mice, suggesting that these changes were too modest to detect at a whole bacterial community level.

As previously observed, there were differences in virome composition between the healthy and UC fecal VLP and bacterial stocks ^7, 37–40^. However, contrasting with previous metagenomic analyses of IBD viromes, there was not a high relative abundance of dsDNA *Caudovirales* phages in our pooled UC VLP stock. Instead, the UC virome was dominated by a single 6,339 bp scaffold classified as *Microviridae*. Our limited sample size to prepare the pooled phage stocks, the known high inter-individual differences in gut viromes ^30^, differences in virome composition based on viral enrichment and DNA extraction strategies ^28, 72, 73^, and the known bias in the amplification of ssDNA viruses using multiple-displacement amplification (MDA) ^74, 75^ are all possible causes that could explain these differences between the virome composition in our dataset and previously published viromes of IBD patients. Still, our observations of increased proportions of temperate phages in the UC VLP stock, and alterations in Bacteroidetes, Actinobacteria, and Proteobacteria in the UC bacterial stock are consistent with previously reported UC-specific alterations of the microbiota ^37, 45–47^. While our strategy to pool the healthy and UC bacterial and VLP stocks comes with the loss of inter-individual genomic information between samples within a given treatment group, pooling stocks maintains intra-specific information while minimizing variation during transplantation, especially important given the stochasticity during microbiota acquisition ^76^ and low percentage of retained taxa following transplantation ^77^. Importantly, we assessed the composition of the pooled viral and bacterial stocks in addition to following changes in microbiota of HMA mice over time, thus allowing us to assess the efficacy of bacterial and VLP engraftment^78^. Additionally, despite the limitations of this approach, compared to previous studies that have investigated VLP-mediated changes to bacterial diversity or disease severity in mice ^21, 23, 24^, our use of HMA mice and human-derived VLPs adds translatability of our findings to the microbiota alterations observed in UC patients.

Importantly, we observed differences in viral beta-diversity post-VLP gavage and identified scaffolds transferred from the pooled VLP stocks to HMA mice, suggesting that phages within the healthy and UC stocks were able to successfully engraft in recipient mice. While VLP shotgun sequencing could not be performed during DSS-colitis, differences in viral diversity persisted into the DSS washout period, suggesting that viromes of mice given healthy and UC VLPs were divergent before and after inflammation. Successful engraftment was also supported by increased viral abundance after VLP gavage and a significant decrease in Jaccard distance of VCs to the healthy stock virome following healthy VLP gavage. While the decrease in Jaccard distance of VCs to the UC stock virome was not significant, these data may be explained by the fact that phages present before VLP gavage in HMA mice colonized by UC bacteria may already resemble the UC stock since they all come from the same initial UC patients. In addition, the post-gavage viromes of healthy and UC VLP-treated were characterized by distinct phage-host interactions, as predicted by CRISPR spacer matches. The treatment-specific differences in the predicted bacterial hosts that these phages target are in support our observations that healthy and UC VLPs have divergent effects on gut bacterial communities. The particular observation that healthy VLPs successfully engrafted in HMA mice colonized with UC bacterial communities is consistent with previous observations of phage “cross-reactivity” between different individuals or disease states and likely supports the idea that some phages in the gut have host-ranges at the species rather than the strain level ^21, 23, 24, 79, 80^. Notably, our functional annotation of viral scaffolds revealed no obvious differences in the auxiliary metabolic genes present between treatment groups (data not shown), suggesting shared potential functionality between viromes.

Regardless of whether HMA mice were given a single dose of healthy or UC VLPs, mice colonized with fecal bacteria isolated from UC patients had increased colitis severity compared to mice colonized with fecal bacteria from healthy donors. This finding is consistent with previous reports showing exacerbation of DSS-induced and T cell-mediated colitis following administration of gut microbiota from IBD patients to germ-free mice ^50, 51^. While these studies associated an increase of IL-17 producing CD4+ T cells to disease severity, anti-viral immunity is more commonly associated with type 1 immunity and interferon production. As such, an impact of the virome on T cell immunity is likely indirect via changes to bacteria-derived products that promote Th17 cell differentiation. Nevertheless, phages express pathogen-associated molecular patterns that, if accessed by host immune cells, may directly alter tissue inflammation ^81^. Although we administered UC VLPs to GF mice in order to investigate this scenario, VLPs were cleared in the absence of bacterial hosts, resulting in undetectable changes to the host immune response.

Two important limitations in using HMA mice to monitor disease outcomes are that (1) only a fraction of human fecal bacteria can colonize GF mice ^77^ and (2) there is a reported colonization bias towards Bacteroidetes ^77^. Despite these limitations, we identified differences in the relative abundances of certain bacterial species in mice after colonization with healthy or UC fecal bacteria, which could explain the differences in colitis severity between these two groups.

This included *Akkermansia* sp., which are reduced in IBD patients and have been shown to be protective in the context of DSS-colitis ^12^, and *Escherichia-Shigella* sp., which are thought to exacerbate the pro-inflammatory immune responses in IBD ^82^. We speculate that the increased colitis severity observed in UC-HMA mice given UC VLPs compared to mice given PBS, heat- killed UC VLPs, or healthy VLPs may be due to alterations of the gut bacterial communities resulting from phage predation. Some of the bacterial species reduced in relative abundance by UC VLPs, such as *E. limosum* ^58^ and *Ruminoclostridium 5,* are thought to be producers of short- chain fatty acids (SCFAs), which are important in promoting epithelial barrier function, antimicrobial peptide production, and induction of immunomodulatory Tregs ^3, 9^. Given that IBD patients have reduced fecal SCFA concentrations ^83, 84^, these data support the idea that UC VLPs may cause increased colitis severity by depleting these SCFA-producing bacteria. In support of this idea, we found that the reduction of *E. limosum* during the DSS/washout period in UC VLP- treated mice coincided with an enrichment of phages predicted to infect *Eubacterium*. However, given the current limitations in bioinformatically matching phages to their hosts (we were only able to assign predicted hosts to ∼ 30% of our viral scaffolds) and the diverse interbacterial interactions in the gut, it is difficult to definitively attribute the phage predation of specific bacterial taxa to an altered disease pathology. Notably, UC VLP administration also led to an increase in the relative abundance of *Escherichia*-*Shigella* sp., possibly an indirect result of phage predation and its subsequent modulation of the gut microbiota through interbacterial interactions, as recently described by Hsu et al. ^19^. Consistent with the notion that Proteobacteria can contribute to IBD pathogenesis ^13^, our data indicate that phage regulation of bacterial communities promotes the expansion of immune-activating, potentially pathogenic bacteria. In agreement with our earlier observations, Khan Mirzaei et al. ^80^ showed that phages from stunted children promoted the *in vitro* expansion of Proteobacteria, which are important in contributing to the nutritional deficiencies of this disease. The observation that VLPs isolated from the patients of two distinct diseases allowed for the expansion of Proteobacteria across different ages and experimental settings has important relevance for virome alterations in inflammatory diseases and warrants further investigation. While it is possible that endotoxin remaining in the VLP preps also contributed to the observed differences in colitis severity, mice given heat-killed UC VLPs displayed reduced colitis severity compared to intact UC VLPs, suggesting that LPS contamination does not explain these effects.

### CONCLUSIONS

While several studies have outlined gut virome alterations in IBD patients, our observations that fecal VLPs from UC patients exacerbate DSS-colitis severity suggest that these alterations could be important for IBD pathogenesis and gut inflammation. This action may be either through phage-mediated changes in the microbiota or by direct interactions with the intestinal immune system. Overall, the power of experimental *in-vivo* cross infections performed here allowed us to highlight a causal role for phages in modulating gut bacterial communities and disease outcome.

## METHODS

### Lead Contact

Further information and requests for resources and reagents should be directed to and will be fulfilled by the lead contact, Corinne Maurice.

### Subject details and sample collection

For all experiments, fecal samples were collected from 3 UC patients (mean age: 38.66 ± 12.97 SD, mean BMI: 28.89 ± 8.14 SD) in remission and 3 unrelated healthy controls (mean age: 42.33 ± 13.65 SD, mean BMI: 23.40 ± 3.91 SD). Samples were stored at -80 °C until processed. Participants of this study were only included if they were older than 18 years and had not taken antibiotics in the 3 months prior to sample collection. UC patients were excluded if they were administered treatments other than immunosuppressants. Human studies were performed with approval of the McGill Ethics Research Board (REB #A04-M27-15B).

### Mice

Six- to 12-week-old female and male germ-free (GF) C57BL/6 mice were purchased from Charles River Laboratories (Wilmington, North Carolina) and maintained in flexible film isolators at McGill University. All mice had unlimited access to autoclaved mouse breeder’s diet and water. All mouse experiments were carried out in accordance with the approved McGill University animal use protocol (7977).

### Experimental model

HMA mice were used to assess how VLP administration impacts the gut bacterial communities and colitis severity. A schematic of each experiment performed, and associated timelines are summarized in Fig. 1. Bacterial communities derived from the feces of UC patients or healthy controls were administered to GF mice by oral gavage (200 uL, 1-3 x 10^8^ cells/mL). All colonized HMA mice were maintained on the same sterilized diet and water as before colonization. During the 3-week bacterial reconstitution period, random fecal samples were weighed and collected from each individual cage every week to monitor the reconstitution. Stool DNA was extracted by using the QIAGEN QIAamp DNA stool mini Kit under a sterile biosafety cabinet. Real-time PCR amplification of the V6 region of the 16S rRNA gene was performed and the concentration of purified PCR products were quantified by NanoDrop spectrophotometer as a standard template. (forward primer: aggattagataccctggta and reverse primer: rrcacgagctgacgac). The load of bacterial DNA in feces was estimated by 16S rRNA gene qPCR, and the concentration was calculated according to the dilution of series of template, and normalized to the stool weight. Following a period of 3 weeks of bacterial reconstitution, 1 or 4 doses of VLPs derived from the feces of UC patients or healthy controls were administered to the HMA mice by oral gavage at equivalent concentrations to the bacterial dose given ^86^ over a 9-10 day period. Controls consisted of the administration of PBS or heat-inactivated VLPs to HMA mice following bacterial reconstitution. Following VLP treatment, 2% DSS was administered to the mice for 5 days, followed by a 5-day washout period. All mice were sacrificed at the end of the 5-day DSS washout period, and mouse colons were removed to determine DSS-colitis severity. Throughout the course of the experiments, mouse fecal pellets were collected to determine bacterial and VLP abundance, bacterial activity and damage, to monitor disease activity index, and to determine bacterial VLP community composition.

### Processing of human fecal samples

VLPs and bacterial communities were extracted and processed from human fecal samples, as described by Khan Mirzaei et al. ^80^ with some modifications. To acquire community-level information and obtain enough material for multiple mouse experiments, two grams of each frozen healthy or UC fecal sample were pooled separately, thawed under anaerobic conditions, resuspended with reduced PBS (rPBS), containing 1 ug.mL^-1^ resazurin sodium salt and 1 mg.mL^-1^ L-Cysteine; final concentrations, thoroughly vortexed, and centrifuged at 800*g* for 1 min at 22°C to remove large debris. Pooling samples still allows for intraspecific genomic information of bacterial and viral communities, despite a loss of inter-individual differences ^87^. The resulting supernatant was centrifuged at 7,000*g* at 22°C for 1 hr to separate bacterial and VLP communities. Following resuspension in rPBS, bacterial concentration was determined using flow cytometry (see below) ^56^. The VLP-containing supernatant was filtered through a 0.2 μm sterile syringe filter (Millex-GP, Millipore Sigma, USA) concentrated by centrifugation (35,000*g,* 4°C) for 3 hr and resuspended in SM buffer. VLP concentration was determined using epifluorescence microscopy (see below) ^22^. DNA extractions were performed on aliquots of each pooled VLP and bacterial preparation for downstream sequencing analysis (see below). As a control, one aliquot of the extracted VLPs were heat-inactivated by incubating the solution at 95°C for 20 min, followed by DNAse (Ambion DNAse I, Thermo-Fisher Scientific, USA) treatment for 2 hr ^22^. Bacterial and viral abundances were confirmed prior to each gavage, and two hundred microliters of bacterial and VLP communities were administered to mice by oral gavage at equal concentrations (1-3 x 10^8^ VLPs or bacterial cells/mL).

### Induction of DSS colitis and disease activity index (DAI) evaluation

HMA and GF mice received 2% (w/v) DSS (MP Biomedicals) in drinking water for 5 days prior to regular drinking water for another 5 days. The amount of DSS intake per mouse was recorded. Each individual mouse was weighed to determine percentage weight changes. Fecal samples were taken from each cage to clinically monitor for rectal bleeding and diarrhea. Hemoccult SENSA kit (Beckman Coulter) was used to assess rectal bleeding as per the manufacturer’s instructions. All parameters were evaluated every 3 days.

### Histological evaluation of colitis

HMA mice were anesthetized and sacrificed five days after completing DSS administration. 0.5 cm of the distal colon was excised, rinsed with saline solution, fixed in 10% formalin and embedded in paraffin. Sections of 4 μm were stained with H&E by the Goodman Cancer Research Centre Histology Facility and assessed for histological changes in a blinded manner.

### Pro-inflammatory cytokine ELISA

The top 0.5 cm of the colon was harvested and cut longitudinally, washed in PBS and excess liquid was removed. The tissue was then weighed, placed in 400 μL of culture medium RPMI1640 supplemented with 10% fetal bovine serum and culture in 37°C incubator overnight. The supernatant was collected and transferred into a 1.5 mL tube and centrifuged for 5 min at 13,000 rpm for 4 °C. The supernatant was collected, stored at −20 °C, and used for ELISA. The mouse IL-6, IL-1β and TNF-α ELISA Kits from Invitrogen (ThermoFisher) were used as per the manufacturer’s instructions.

### Colon cell extraction

The middle 5 cm of the colon was used to generate single cell suspensions of lamina propria cells. The tissue was cut longitudinally and washed in cold Hank’s Balanced Salt Solution (HBSS) + EDTA buffer. The tissue was then cut into 0.5–1 cm sections, placed in HBSS + EDTA buffer and incubated at 37 °C for two 20 min periods with washing between incubations. The tissue was then washed twice with cold HBSS buffer and digested in 5mL of digestion buffer (RPMI 1640 supplemented with 10% FBS, 200 U/mL of Collagenase VIII) for 25 min at 37 °C. After digestion, the tissue was passed through a 100 μm filter and resuspended in PBS + 5% FBS for counting using Trypan blue and flow cytometry analysis.

### Flow Cytometry of Colon Cells

Colon cell suspensions were incubated with a fixable Viability dye (eFluor 506, eBioscience) for 25 min at 4 °C. Cells were then incubated with Fc block (7 min at 4 °C), followed by staining (for 30 min at 4 °C) with the following antibodies in appropriate combinations of fluorophores. From Invitrogen: CD45.2 (104). From Biolegend: CD11b (M1/70), Ly6c (HK1.4), Ly6g (1A8). Data were acquired with a FACS Canto II or LSR Fortessa (BD Biosciences) and analyzed using FlowJo software (TreeStar).

### Processing mouse fecal samples for microbiota analyses

Mouse fecal samples were processed similar to human fecal samples, with some modifications. On each sampling date, fecal pellets were collected from each mouse and samples belonging to the same cage were pooled in order to maximize the amount of biological material obtained. Pooled pellets were resuspended in rPBS under anaerobic conditions, thoroughly vortexed and centrifuged at 800*g* for 1 min at 22°C to remove large debris. The resulting supernatant was centrifuged at 6,000*g* for 5 min at 22°C to separate VLP and bacterial communities. VLP-containing supernatant was filtered through a 0.2 um sterile syringe filter (Millex-GP, Millipore Sigma, USA) to remove the remaining bacteria, and phage abundance was determined using epifluorescence microscopy (see below). The bacterial pellet was washed twice and resuspended in rPBS under anaerobic conditions. The concentration and proportion of active and damaged bacterial cells were determined using flow cytometry (see below). For the remaining fecal pellets from each cage, bacterial and VLP communities were separated, as described above. Following separation, bacterial and VLP DNA were extracted separately.

### Bacterial DNA extraction and 16S rRNA gene sequencing analyses

Bacterial DNA was extracted from human and mouse feces-derived bacterial supernatant using the DNeasy powersoil kit (Qiagen, USA) according to the manufacturer’s instructions. Data included from healthy and UC bacterial stocks are from four and five technical sequencing replicates, respectively. The V4 region of the 16S rRNA gene was amplified using the 515F/806R primers, and pooled amplicons were sequenced on an Illumina Miseq with 250 bp paired-end technology at the Genome Québec *Centre d’expertise et de services* core facility ^88^. 16S rRNA gene sequencing analysis was performed using the QIIME2 platform (v 2020.2, v2020.6) ^89^. Read- trimming, removal of chimeric reads, merging of paired-end reads, and inference of amplicon sequence variants was performed using DADA2 ^90^. In each sequencing run, read depth was rarefied to the lowest number of reads that a sample contained in that run. Bacterial diversity between treatment groups was assessed using Weighted UniFrac and PcoA. Taxonomic identification was performed using the QIIME2 feature classifier, training a Naives-Bayes Classifier on the Silva 138 database ^91^. The statistical framework, ANCOM, was used to identify differentially abundant bacterial taxa between treatment groups ^49^. In order to disentangle the effects of VLP treatment and possible isolator-specific colonization biases, bacterial species that were found to be differentially abundant during the bacterial colonization period were removed from Supplementary Tables. S5-8. In addition, each differentially abundant bacterial species was assigned a ranking (1-3) based on likelihood that differences in relative abundance were due to VLP treatment or isolator effect (1: likely due to phage treatment, 2: possibly due to phage treatment, 3: likely due to isolator effect). Species classified as “likely due to phage treatment (1)” included species, where reads were detected in a majority of cages in all treatment groups. Species classified as “possibly due to VLP treatment” (2) included species that were found to be differentially abundant in the VLP gavage and/or DSS washout period and where reads were not detected in a majority of cages in all treatment groups during the bacterial colonization period. Any differentially abundant species that did not meet the above criteria were classified as “likely due to isolator effect” (3).

### VLP DNA extraction, amplification and sequencing

VLP enrichment and subsequent DNA extraction was performed as described by Reyes et al. ^22^ with some modifications. Briefly, feces-derived VLP supernatant (prepared as described above) was incubated with lysozyme (45 min at 37 °C, 50 mg·mL^-1^), Ambion™ Dnase I (1 hr at 37 °C, 2U), and proteinase K (1 hr at 37 °C, 20 mg·mL^-1^). After addition of 5M NaCl and 10% Cetyltrimethylammonium Bromide (CTAB)/0.7M NaCl solutions, samples were transferred to phase lock gel tubes (QuantaBio, USA) with an equal amount of phenol:chloroform:isoamyl alcohol (25:24:1 v/v, pH = 8.0, ThermoFisher Scientific, USA). The upper aqueous, DNA-containing, layer was transferred to a new tube and left to precipitate overnight at -80 °C in 100% ice-cold ethanol. Precipitated samples were then purified with the Zymo DNA Clean & Concentrator 25 kit (Zymo Research, USA). Following purification, 1uL of DNA was amplified in triplicate using MDA, using the GenomiPhi V3 DNA amplification kit (Cytiva, USA). Amplified products were pooled, and DNA concentrations were quantified with the Qubit dsDNA high-sensitivity (HS) assay kit (ThermoFisher Scientific, USA). In total, 27/90 samples from HMA mice in the experiment referenced in Fig. 1B (trial #1) were excluded from sequencing due to insufficient DNA yield, including all 18 samples (2 sample collection dates) during the DSS-colitis period. For virome analyses of human stocks, 4 (healthy) and 5 (UC) technical replicates were sequenced from the same pooled fecal samples. Paired-end Illumina shotgun sequencing libraries were prepared from purified VLP-derived DNA at the Genome Québec *Centre d’expertise et de services* core facility. Barcoded libraries which passed quality control were pooled and sequenced on an Illumina MiSeq with 250 bp paired-end sequencing technology.

### VLP Abundances

VLPs were enumerated using epifluorescence microscopy ^33^. Briefly, human or mouse-derived VLP supernatant were diluted in TE buffer and fixed with 1% formaldehyde for 15 min. Fixed VLPs were filtered onto 0.02 μm filter membranes in triplicate (Anodisc, GE Healthcare, USA) and were stained with SYBR-Gold (2.5X concentration). On average, 25 fields of view of the stained filters were visualized and counted in triplicate using an epifluorescence microscope (Zeiss, Axioskop).

### Bacterial Abundance and Physiology

The concentration of bacterial cells and the proportion of active and damaged bacterial cells in human and mouse feces was determined using flow cytometry, as described previously ^56^. Briefly, feces-derived bacterial supernatant was diluted in rPBS anaerobically, stained with SYBRGreen I (1x concentration, 15 min) or PI (1x concentration, 10 min) and counted using flow cytometry on a FACSCanto II (BD, USA) equipped with a 488nm laser (20 mW) and 530/30 and 585/42 detection filters. Five-to-ten microliters of green fluorescent reference beads of 3.0-3.4 μm (Rainbow beads, BD Biosciences) were added to each flow cytometry tube prior to acquisition to calculate bacterial concentration. The concentration of the Rainbow beads was determined by calibration with Trucount tubes (BD, USA) for each sampling date ^92^. Total bacterial abundance was determined by SYBRGreen I staining. As previously described, ^92^ relative nucleic acid content was used as a proxy for bacterial activity: active cells, containing more nucleic acid, were identified by their higher levels of green fluorescence. Bacterial membrane damage was assessed by PI staining: cells with high red fluorescence have lost their membrane integrity. All stainings were done in triplicates. Gating was performed using the FlowJo analysis software (v10.6.1). Gating strategy is shown in Supplementary Fig. S5A-B.

### Prophage induction assay

In order to determine whether DSS could directly cause prophage induction of fecal bacteria, we performed an *in vitro* induction assay ^62^. Briefly, pooled UC fecal bacteria were grown anaerobically in triplicate at 37°C, mixing every 15 min, in reduced Brain Heart Infusion broth (BBL BD, Mississauga, ON, Canada) supplemented with hemin (5 μg/mL) and vitamin K (1 μg/mL). At early exponential phase, DSS was added to the growth media at final concentrations of 0% (H_2_O added), 1%, 2%, or 5%. Bacterial growth was monitored every 15 min for 14 hr, measuring OD_600_ (Epoch 2 microplate spectrophotometer, Biotek Instruments, Winooski, VT, USA) until stationary phase was reached. Bacteria were pelleted (2,000xg, 15 min, 22°C) and VLP-containing supernatant was fixed with 1% formaldehyde and enumerated using epifluorescence microscopy. Induction was identified through a concomitant significant decrease in bacterial abundance and increase in VLPs.

### Bioinformatic analysis of VLP sequence data

Virome analysis was performed on shotgun-sequenced VLP DNA, as described in Supplementary Fig. S1A. Briefly, raw reads (mean reads mouse samples: 176,955 ± 21,451 SD; mean reads human VLP stocks: 1,176,926 ± 79038 SD) were first trimmed using Trimmomatic (v0.38) ^93^. *De novo* assembly of sequencing reads was carried out with SPAdes ^94^ (v3.12.0), using default settings. Scaffolds from each sample’s assembly were then pooled and those less than 3 kb in length were removed ^30^. These scaffolds were used as input for VirSorter v1.0.6 ^95^ with the following settings: virome decontamination, DIAMOND, viromes + Gut Virome Database v1.7.2018 ^28^. VirSorter- positive scaffolds were then used as input for VIBRANT (v1.2.1) ^29^ with the virome mode enabled. Scaffolds which were classified as viral by VirSorter and then subsequently by VIBRANT were retained for downstream taxonomic and diversity analysis. Scaffolds classified as viral from mouse samples and scaffolds classified as viral from the pooled human stock samples were pooled, and scaffolds with 90% nucleotide identity over 90% of the shorter scaffold’s length were removed using CD-HIT-EST v4.8.1 ^96^ to form the non-redundant scaffolds library. vConTACT2 v0.9.10 was used to form viral clusters from viral scaffolds ^44^. These clusters were added to the scaffold metadata used for downstream analysis. Demovir (https://github.com/feargalr/Demovir) was used to assign taxonomic identities to the viral scaffolds. This tool uses a voting approach for taxonomy based on protein homology to a viral subset of the TrEMBL database from UniProt. Additionally, scaffolds were identified as being a crAss-like phage if they had BLASTn alignments to a crAssphage database with an e-value ≤ 10^-10^, covering ≥ 90% of the scaffold’s length, and over 50% nucleotide identity ^30, 97^. We used CrisprOpenDB, which uses BLASTn alignment to a CRISPR spacer database to assign bacterial genus predictions for each viral scaffold ^48^. Default settings were used, allowing up to 2 mismatches between a scaffold and spacer. Using VIBRANT, scaffolds were identified as temperate based on the presence of the integrase gene ^29^. We used Bowtie2 v2.3.5 ^98^ to map trimmed reads back to assembled viral scaffolds. SAMtools v1.9 ^99^ was used to convert SAM to BAM files. Custom Python scripts were used to create scaffold metadata files and abundance matrices. These scripts are available at the GitHub (https://github.com/MauriceLab/phage_colitis). Scaffolds were only included in analyses for a given sample if coverage was ≥ 1x over ≥75% of the scaffold length ^100^. In order to determine differentially abundant viral scaffolds, raw counts matrices were input into DESeq2 v.1.3.0 ^57^. The standard DESeq2 workflow was performed, using “poscounts” for estimation of size factors. In each comparison, scaffolds with adjusted p-values < 0.001 and with log_2_fold changes greater or less than 1 were considered differentially abundant ^85^. Bray-Curtis dissimilarity and Jaccard distances for diversity analyses were performed on matrices normalized by total coverage using Vegan v2.5-7. Visualization of NMDS and relative abundance plots was performed using the ggplot2 v3.3.3 package.

### Quantification and Statistical Analysis

Data were analyzed using GraphPad Prism software (v 8.4.3). Specific tests for determining statistical significance are indicated in the figure legends. P values < 0.05 were considered statistically significant, except for DESeq2 virome analyses, where p values < 0.001 and values with log_2_fold change greater or less than 1 were considered statistically significant. Differences in viral diversity were assessed using NMDS of Bray-Curtis dissimilarity and Jaccard distance in R v.4.0.5 (R Core Team, 2021) using the Vegan v2.5-7 and ggplot2 v3.3.3 packages. Adonis PERMANOVA was used to determine significant differences in Bray-Curtis Dissimilarity between the viromes of HMA mice given healthy VLPs, UC VLPs or PBS, using the Vegan v2.5- 7 package. Differentially abundant bacterial species were identified using ANCOM. PcoA on weighted UniFrac distances were used to determine differences in bacterial diversity between HMA mice. Significant differences on weighted UniFrac distance between HMA mice was determined using PERMANOVA, using the QIIME2 platform (v2020.2, v2020.6) ^89^. P-values for all PERMANOVA tests were adjusted for multiple comparisons using Benjamini-Hochberg.

## DECLARATIONS

### Ethics approval and consent to participate

Human studies were performed with approval of the McGill Ethics Research Board (REB #A04- M27-15B). All participants provided informed consent prior to donating samples. All mouse experiments were carried out in accordance with the approved McGill University animal use protocol (7977).

### Availability of data and materials

Custom scripts and code used for data analysis are available at https://github.com/MauriceLab/phage_colitis. Sequencing reads for bacterial 16S rRNA gene sequencing and VLP shotgun metagenomics are available on NCBI SRA. Bacterial 16S rRNA gene sequencing reads are available using accession number: PRJNA734661. VLP shotgun metagenomic reads are available using accession number: PRJNA732769. This study did not generate new unique reagents.

### Authors’ Contributions

A.S., Y.L. and M.K.M. designed and performed experiments, analyzed results and wrote the manuscript. A.S. and M.S. performed bioinformatic analyses. R.S. provided critical animal resources. C.F.M. and I.L.K. conceived the project, designed experiments and wrote the manuscript.

## Acknowledgements

A special thanks to the Comparative Medicine and Animal Resources Centre of McGill University, especially Anna Jimenez for her technical support using germ-free mice. We also appreciate the support of Genome Quebec for microbial sequencing services. The flow cytometry work was performed in the Flow Cytometry Core Facility of the Life Science Complex and supported by funding from the Canadian Foundation for Innovation. AS is supported by a Canadian Institute of Health Research (CIHR) fellowship (170921). YL was supported by a Richard and Edith Strauss Postdoctoral Fellowship in Medicine. ILK is a Canada Research Chair in Barrier Immunity. CFM is a Canada Research Chair in Gut Microbial Physiology and CIFAR Azrieli Global Scholar, Humans & Microbiome Program.

## ADDITIONAL FILES

### Additional File 1

**Supplementary Figure S1.** Virome analyses on stock and mouse VLPs. Related to Figure 2 and Figure 5. **Supplementary Figure S2.** PcoA on Weighted UniFrac distances in HMA mice given VLPs. Related to Figure 3 and Figure 6. **Supplementary Figure S3.** Experimental colitis severity of UC-HMA and healthy-HMA mice given a single dose of healthy or UC VLPs. Related to Figure 3. **Supplementary Figure S4.** Viral and bacterial abundance in UC-HMA mice. Related to Figure 4. **Supplementary Figure S5.** Bacterial physiology of HMA mice. Related to Figure 4. **Supplementary Figure S6.** Human microbiota protects mice from experimental colitis. Related to Figure 7. **Supplementary Table. S2.** PERMANOVA on weighted UniFrac distances between UC or healthy-HMA mice given healthy VLPs or UC VLPs. Related to Figure 3 and Figure 6. **Supplementary Table. S5.** Differentially abundant species during VLP gavage period in HMA mice given healthy VLPs, UC VLPs, or PBS. Related to Figure 6. **Supplementary Table. S6.** Differentially abundant species during DSS/washout period in HMA mice given healthy VLPs, UC VLPs, or PBS. Related to Figure 6. **Supplementary Table. S7.** Differentially abundant species during VLP gavage period in HMA mice given UC VLPs (+/- DSS), or heat-killed UC VLPs. Related to Figure 6. **Supplementary Table. S8.** Differentially abundant species during DSS/ washout period in HMA mice given UC VLPs (+/- DSS) or heat-killed UC VLPs. Related to Figure 6.

### Additional File 2

**Table. S1.** VLP scaffolds and bacterial phyla and genera in pooled healthy and UC stocks. Related to Figure 2. Healthy VLP stock scaffolds and annotations (tab 1), UC VLP stock scaffolds and annotations (tab 2), bacterial phyla relative abundance (tab 3), bacterial genera relative abundance (tab 4).

### Additional File 3

**Table. S3.** VLP scaffolds shared between stocks and HMA mice post-VLP gavage. Related to Figure 5. VLP scaffolds shared between healthy VLP stock and UC-HMA mice post-VLP gavage (tab 1), VLP scaffolds shared between the UC VLP stock and UC-HMA mice post-VLP gavage (tab 2), scaffold metadata (tab 3).

### Additional File 4

**Table. S4.** Differentially abundant viral scaffolds. Related to Figure 5. Healthy VLPs vs. PBS control, phage gavage period (tab 1). UC VLPs vs. PBS control, phage gavage period (tab 2). Healthy VLPs vs. PBS control, DSS washout period (tab 3). UC VLPs vs. PBS control, DSS washout period (tab 4). Post-DSS vs. pre-DSS, PBS control (tab 5). Post-DSS vs. pre-DSS, HMA- mice treated with healthy VLPs (tab 6). Post-DSS vs. pre-DSS, HMA mice treated with UC VLPs (tab 7).

## REFERENCES

1. Valdes, A. M., Walter, J., Segal, E. & Spector, T. D. Role of the gut microbiota in nutrition and health. Bmj 361, k2179, doi:10.1136/bmj.k2179 (2018).

2. Belkaid, Y. & Hand, T. W. Role of the microbiota in immunity and inflammation. Cell 157, 121–141, doi:10.1016/j.cell.2014.03.011 (2014).

3. Arpaia, N. et al. Metabolites produced by commensal bacteria promote peripheral regulatory T-cell generation. Nature 504, 451–455, doi:10.1038/nature12726 (2013).

4. Atarashi, K. et al. Treg induction by a rationally selected mixture of Clostridia strains from the human microbiota. Nature 500, 232–236, doi:10.1038/nature12331 (2013).

5. Furusawa, Y. et al. Commensal microbe-derived butyrate induces the differentiation of colonic regulatory T cells. Nature 504, 446–450, doi:10.1038/nature12721 (2013).

6. Pabst, O. et al. Adaptation of solitary intestinal lymphoid tissue in response to microbiota and chemokine receptor CCR7 signaling. J Immunol 177, 6824–6832, doi:doi:10.4049/jimmunol.177.10.6824 (2006).

7. Lloyd-Price, J. et al. Multi-omics of the gut microbial ecosystem in inflammatory bowel diseases. Nature 569, 655–662, doi:https://doi.org/10.1038/s41586-019-1237-9 (2019).

8. Nemoto, H. et al. Reduced diversity and imbalance of fecal microbiota in patients with ulcerative colitis. Dig Dis Sci 57, 2955–2964, doi:10.1007/s10620-012-2236-y (2012).

9. Parada Venegas, D. et al. Short Chain Fatty Acids (SCFAs)-Mediated Gut Epithelial and Immune Regulation and Its Relevance for Inflammatory Bowel Diseases. Frontiers of Immunology 10, 277, doi:10.3389/fimmu.2019.00277 (2019).

10. Rigottier-Gois, L. Dysbiosis in inflammatory bowel diseases: the oxygen hypothesis. ISME 7, 1256–1261, doi:10.1038/ismej.2013.80 (2013).

11. Bel, S. et al. Reprogrammed and transmissible intestinal microbiota confer diminished susceptibility to induced colitis in TMF −/− mice. PNAS 13, 4964–4969, doi:doi: 10.1073/pnas.1319114111 (2014).

12. Bian, X. et al. Administration of Akkermansia muciniphila Ameliorates Dextran Sulfate Sodium-Induced Ulcerative Colitis in Mice. Frontiers in Microbiology 10, doi:10.3389/fmicb.2019.02259 (2019).

13. Garrett, W. S. et al. Enterobacteriaceae act in concert with the gut microbiota to induce spontaneous and maternally transmitted colitis. Cell Host Microbe 8, 292–300, doi:10.1016/j.chom.2010.08.004 (2010).

14. Lee, J. Y. et al. High-Fat Diet and Antibiotics Cooperatively Impair Mitochondrial Bioenergetics to Trigger Dysbiosis that Exacerbates Pre-inflammatory Bowel Disease. Cell Host Microbe 28, 273–284.e276, doi:10.1016/j.chom.2020.06.001 (2020).

15. Martín, R. et al. Faecalibacterium prausnitzii prevents physiological damages in a chronic low-grade inflammation murine model. BMC Microbiology 15, 67, doi:doi:10.1186/s12866-015-0400-1 (2015).

16. Graham, D. B. & Xavier, R. J. Pathway paradigms revealed from the genetics of inflammatory bowel disease. Nature 578, 527–539, doi:10.1038/s41586-020-2025-2 (2020).

17. Jostins, L. et al. Host–microbe interactions have shaped the genetic architecture of inflammatory bowel disease. Nature 491, 119–124, doi:10.1038/nature11582 (2012).

18. Breitbart, M., Bonnain, C., Malki, K. & Sawaya, N. A. Phage puppet masters of the marine microbial realm. Nature Microbiology 3, 754–766, doi:doi: 10.1038/s41564-018-0166-y (2018).

19. Hsu, B. et al. Dynamic Modulation of the Gut Microbiota and Metabolome by Bacteriophages in a Mouse Model. Cell Host and Microbe 6, 803–815 doi:https://doi.org/10.1101/454579 (2018).

20. Kim, M. S., Park, E. J., Roh, S. W. & Bae, J. W. Diversity and abundance of single- stranded DNA viruses in human feces. Appl Environ Microbiol 77, 8062–8070, doi:10.1128/aem.06331-11 (2011).

21. Rasmussen, T. S. et al. Faecal virome transplantation decreases symptoms of type 2 diabetes and obesity in a murine model. Gut 69, 2122–2130, doi:10.1136/gutjnl-2019-320005 (2020).

22. Reyes, A., Wu, M., McNulty, N. P., Rohwer, F. L. & Gordon, J. I. Gnotobiotic mouse model of phage-bacterial host dynamics in the human gut. Proc Natl Acad Sci U S A 110, 20236–20241, doi:10.1073/pnas.1319470110 (2013).

23. Draper, L. A. et al. Autochthonous faecal viral transfer (FVT) impacts the murine microbiome after antibiotic perturbation. BMC Biology 18, 1–14 (2020).

24. Lin, D. M. et al. Transplanting Fecal Virus-Like Particles Reduces High-Fat Diet- Induced Small Intestinal Bacterial Overgrowth in Mice. Front Cell Infect Microbiol 9, 348, doi:10.3389/fcimb.2019.00348 (2019).

25. Ott, S. J. et al. Efficacy of Sterile Fecal Filtrate Transfer for Treating Patients With Clostridium difficile Infection. Gastroenterology 152, 799–811.e797, doi:10.1053/j.gastro.2016.11.010 (2017).

26. Khan Mirzaei, M. & Maurice, C. F. Ménage à trois in the human gut: interactions between host, bacteria and phages. Nat Rev Microbiol 15, 397–408, doi:10.1038/nrmicro.2017.30 (2017).

27. Camarillo-Guerrero, L. F., Almeida, A., Rangel-Pineros, G., Finn, R. D. & Lawley, T. D. Massive expansion of human gut bacteriophage diversity. Cell 184, 1098–1109.e1099, doi:10.1016/j.cell.2021.01.029 (2021).

28. Gregory, A. C. et al. The Gut Virome Database Reveals Age-Dependent Patterns of Virome Diversity in the Human Gut. Cell Host Microbe 28, 724–740.e728, doi:10.1016/j.chom.2020.08.003 (2020).

29. Kieft, K., Zhou, Z. & Anantharaman, K. VIBRANT: automated recovery, annotation and curation of microbial viruses, and evaluation of viral community function from genomic sequences. Microbiome 8 (2020).

30. Shkoporov, A. N. et al. The Human Gut Virome Is Highly Diverse, Stable, and Individual Specific. Cell Host and Microbe 26, 527–541 (2019).

31. Minot, S. et al. Rapid evolution of the human gut virome. Proc Natl Acad Sci U S A 110, 12450–12455, doi:10.1073/pnas.1300833110 (2013).

32. Fujimoto, K. et al. Metagenome Data on Intestinal Phage-Bacteria Associations Aids the Development of Phage Therapy against Pathobionts. Cell Host Microbe 28, 380–389.e389, doi:10.1016/j.chom.2020.06.005 (2020).

33. Reyes, A. et al. Viruses in the faecal microbiota of monozygotic twins and their mothers. Nature 466, 334–338, doi:10.1038/nature09199 (2010).

34. Zuo, T. et al. Human-Gut-DNA Virome Variations across Geography, Ethnicity, and Urbanization. Cell Host Microbe 28, 741–751.e744, doi:10.1016/j.chom.2020.08.005 (2020).

35. Kim, M.-S. & Bae, J.-W. Lysogeny is prevalent and widely distributed in the murine gut microbiota. ISME 12, 1127 – 1141, doi:10.1038/s41396-018-0061-9 (2018).

36. Edwards, R. A. et al. Global phylogeography and ancient evolution of the widespread human gut virus crAssphage. Nat Microbiol 4, 1727–1736, doi:10.1038/s41564-019-0494-6 (2019).

37. Clooney, A. G. et al. Whole-Virome Analysis Sheds Light on Viral Dark Matter in Inflammatory Bowel Disease. Cell Host Microbe 26, 764–778.e765, doi:10.1016/j.chom.2019.10.009 (2019).

38. Fernandes, M. A. et al. Enteric Virome and Bacterial Microbiota in Children With Ulcerative Colitis and Crohn Disease. J Pediatr Gastroenterol Nutr 68, 30–36, doi:10.1097/mpg.0000000000002140 (2019).

39. Norman, J. M. et al. Disease-specific alterations in the enteric virome in inflammatory bowel disease. Cell 160, 447–460, doi:10.1016/j.cell.2015.01.002 (2015).

40. Zuo, T. et al. Gut mucosal virome alterations in ulcerative colitis. Gut 68, doi:doi: 10.1136/gutjnl-2018-318131). (2019).

41. Cornuault, J. K. et al. Phages infecting Faecalibacterium prausnitzii belong to novel viral genera that help to decipher intestinal viromes. Microbiome 6, 65, doi:10.1186/s40168-018-0452-1 (2018).

42. Duerkop, B. A. et al. Murine colitis reveals a disease-associated bacteriophage community. Nat Microbiol 3, 1023–1031, doi:10.1038/s41564-018-0210-y (2018).

43. Okayasu, I. et al. A novel method in the induction of reliable experimental acute and chronic ulcerative colitis in mice. Gastroenterology 98, 694–702, doi:10.1016/0016-5085(90)90290-h (1990).

44. Bin Jang, H. et al. Taxonomic assignment of uncultivated prokaryotic virus genomes is enabled by gene-sharing networks. Nat Biotechnol 37, 632–639, doi:10.1038/s41587-019-0100-8 (2019).

45. Zhou, Y. & Zhi, F. Lower Level of Bacteroides in the Gut Microbiota Is Associated with Inflammatory Bowel Disease: A Meta-Analysis. Biomed Res Int 2016, 5828959, doi:10.1155/2016/5828959 (2016).

46. Alam, M. T. et al. Microbial imbalance in inflammatory bowel disease patients at different taxonomic levels. Gut Pathog 12, 1, doi:10.1186/s13099-019-0341-6 (2020).

47. Zakerska-Banaszak, O. et al. Dysbiosis of gut microbiota in Polish patients with ulcerative colitis: a pilot study. Sci Rep 11 (2021).

48. Dion, M. B. et al. Streamlining CRISPR spacer-based bacterial host predictions to decipher the viral dark matter. Nucleic Acids Res 49, 3127–3138, doi:10.1093/nar/gkab133 (2021).

49. Mandal, S. et al. Analysis of composition of microbiomes: a novel method for studying microbial composition. Microb Ecol Health Dis 29, doi:doi: 10.3402/mehd.v26.27663 (2015).

50. Britton, G. J. et al. Microbiotas from Humans with Inflammatory Bowel Disease Alter the Balance of Gut Th17 and RORγt(+) Regulatory T Cells and Exacerbate Colitis in Mice. Immunity 50, 212–224.e214, doi:10.1016/j.immuni.2018.12.015 (2019).

51. Natividad, J. M. et al. Ecobiotherapy Rich in Firmicutes Decreases Susceptibility to Colitis in a Humanized Gnotobiotic Mouse Model. Inflamm Bowel Dis 21, 1883–1893, doi:10.1097/mib.0000000000000422 (2015).

52. Png, C. W. et al. Mucolytic bacteria with increased prevalence in IBD mucosa augment in vitro utilization of mucin by other bacteria. Am J Gastroenterol 105, 2420–2428, doi:10.1038/ajg.2010.281 (2010).

53. Abedon, S. T., Kuhl, S. J., Blasdel, B. G. & Kutter, E. M. Phage treatment of human infections. Bacteriophage 1, 66–85, doi:10.4161/bact.1.2.15845 (2011).

54. Bull, J. J., Vegge, C. S., Schmerer, M., Chaudhry, W. N. & Levin, B. R. Phenotypic resistance and the dynamics of bacterial escape from phage control. PLoS One 9, e94690, doi:10.1371/journal.pone.0094690 (2014).

55. Silveira, C. B. & Rohwer, F. L. in NPJ Biofilms Microbiomes Vol. 2 16010 (2016).

56. Maurice, C. F., Haiser, H. J. & Turnbaugh, P. J. Xenobiotics shape the physiology and gene expression of the active human gut microbiome. Cell 152, 39–50, doi:10.1016/j.cell.2012.10.052 (2013).

57. Love, M. I., Huber, W. & Anders, S. Moderated estimation of fold change and dispersion for RNA-seq data with DESeq2. Genome Biology 15, doi:10.1186/s13059-014-0550-8 (2014).

58. Kanauchi, O. et al. Eubacterium limosum ameliorates experimental colitis and metabolite of microbe attenuates colonic inflammatory action with increase of mucosal integrity. World J Gastroenterol 12, 1071–1077, doi:10.3748/wjg.v12.i7.1071 (2006).

59. Takahashi, K. et al. Effect of Enterococcus faecalis 2001 on colitis and depressive-like behavior in dextran sulfate sodium-treated mice: involvement of the brain–gut axis. Journal of Neuroinflammation 16 (2019).

60. Seishima, J. et al. Gut-derived Enterococcus faecium from ulcerative colitis patients promotes colitis in a genetically susceptible mouse host. Genome Biol 20, 252, doi:10.1186/s13059-019-1879-9 (2019).

61. Frank, D. N. et al. Molecular-phylogenetic characterization of microbial community imbalances in human inflammatory bowel diseases. Proc Natl Acad Sci U S A 104, 13780–13785, doi:10.1073/pnas.0706625104 (2007).

62. Sutcliffe, S. G., Shamash, M., Hynes, A. P. & Maurice, C. F. Common Oral Medications Lead to Prophage Induction in Bacterial Isolates from the Human Gut. Viruses 13, 455 (2021).

63. Otsuji, N., Sekiguchi, M., Iijima, T. & Takagi, Y. Induction of phage formation in the lysogenic Escherichia coli K-12 by mitomycin C. Nature 184(**Suppl 14**), 1079–1080, doi:10.1038/1841079b0 (1959).

64. Jiang, S. C. & Paul, J. H. Occurrence of lysogenic bacteria in marine microbial communities as determined by prophage induction. Mar Ecol Prog Ser 142, 27–38 (1996).

65. Braga, L. P. P. et al. Impact of phages on soil bacterial communities and nitrogen availability under different assembly scenarios. Microbiome 8, doi:doi: 10.1186/s40168-020-00822-z (2020).

66. Shkoporov, A. N. & Hill, C. Bacteriophages of the Human Gut: The “Known Unknown” of the Microbiome. Cell Host Microbe 25, 195–209, doi:10.1016/j.chom.2019.01.017 (2019).

67. Gogokhia, L. et al. Expansion of Bacteriophages Is Linked to Aggravated Intestinal Inflammation and Colitis. Cell Host Microbe 25, 285–299.e288, doi:10.1016/j.chom.2019.01.008 (2019).

68. Schirmer, M. et al. Dynamics of metatranscription in the inflammatory bowel disease gut microbiome. Nat Microbiol 3, 337–346, doi:10.1038/s41564-017-0089-z (2018).

69. Berry, D. et al. Phylotype-level 16S rRNA analysis reveals new bacterial indicators of health state in acute murine colitis. Isme j 6, 2091–2106, doi:10.1038/ismej.2012.39 (2012).

70. Kiesler, P., Fuss, I. J. & W, S. Experimental Models of Inflammatory Bowel Diseases. Cellular and Molecular Gastroenterology and Hepatology 1, 154–170, doi:https://doi.org/10.1016/j.jcmgh.2015.01.006 (2015).

71. Krych, Ł. et al. Have you tried spermine? A rapid and cost-effective method to eliminate dextran sodium sulfate inhibition of PCR and RT-PCR. Journal of Microbiological Methods 144, 1–7 (2018).

72. Castro-Mejía, J. L. et al. Optimizing protocols for extraction of bacteriophages prior to metagenomic analyses of phage communities in the human gut. Microbiome 3, doi:10.1186/s40168-015-0131-4. (2015).

73. Kleiner, M., Hooper, L. V. & Duerkop, B. A. Evaluation of methods to purify virus-like particles for metagenomic sequencing of intestinal viromes. BMC genomics 16, doi:10.1186/s12864-014-1207-4 (2015).

74. Kim, K. H. & Bae, J. W. Amplification methods bias metagenomic libraries of uncultured single-stranded and double-stranded DNA viruses. Appl Environ Microbiol 77, 7663–7668, doi:10.1128/aem.00289-11 (2011).

75. Parras-Moltó, M., Rodríguez-Galet, A., Suárez-Rodríguez, P. & López-Bueno, A. Evaluation of bias induced by viral enrichment and random amplification protocols in metagenomic surveys of saliva DNA viruses. Microbiome 6, doi:10.1186/s40168-018-0507-3 (2018).

76. McCafferty, J. et al. Stochastic changes over time and not founder effects drive cage effects in microbial community assembly in a mouse model. ISME 7, 2116–2125 (2013).

77. Fouladi, F. et al. Sequence variant analysis reveals poor correlations in microbial taxonomic abundance between humans and mice after gnotobiotic transfer. ISME 14, 1809 – 1820 (2020).

78. Arrieta, M. C., Walter, J. & Finlay, B. B. Human Microbiota-Associated Mice: A Model with Challenges. Cell Host Microbe 19, 575–578, doi:10.1016/j.chom.2016.04.014 (2016).

79. Džunková, M. et al. Defining the human gut host–phage network through single-cell viral tagging. Nature Microbiology 4, 2192–2203 (2019).

80. Khan Mirzaei, M. et al. Bacteriophages Isolated from Stunted Children Can Regulate Gut Bacterial Communities in an Age-Specific Manner. Cell Host Microbe 27, 199–212.e195, doi:10.1016/j.chom.2020.01.004 (2020).

81. Li, Y., Handley, S. A. & Baldridge, M. T. The dark side of the gut: Virome-host interactions in intestinal homeostasis and disease. J Exp Med 218, doi:10.1084/jem.20201044 (2021).

82. Mirsepasi-Lauridsen, H. C., Vallance, B. A., Krogfelt, K. A. & Petersen, A. M. Escherichia coli Pathobionts Associated with Inflammatory Bowel Disease. Clin Microbiol Rev 32, doi:10.1128/cmr.00060-18 (2019).

83. Machiels, K. et al. A decrease of the butyrate-producing species Roseburia hominis and Faecalibacterium prausnitzii defines dysbiosis in patients with ulcerative colitis. Gut 63, 1275–1283, doi:10.1136/gutjnl-2013-304833 (2014).

84. Huda-Faujan, N. et al. The impact of the level of the intestinal short chain Fatty acids in inflammatory bowel disease patients versus healthy subjects. Open Biochem J 4, 53–58, doi:10.2174/1874091x01004010053 (2010).

85. Mangalea, M. R. et al. Individuals at risk for rheumatoid arthritis harbor differential intestinal bacteriophage communities with distinct metabolic potential. Cell Host Microbe 29, 726–739.e725, doi:10.1016/j.chom.2021.03.020 (2021).

86. Staley, C. et al. Stable engraftment of human microbiota into mice with a single oral gavage following antibiotic conditioning. Microbiome 5, doi:doi: 10.1186/s40168-017-0306-2 (2017).

87. Schlötterer, C., Tobler, R., Kofler, R. & Nolte, V. Sequencing pools of individuals — mining genome-wide polymorphism data without big funding. Nat Rev Genet 15, 749– 763 (2014).

88. Caporaso, J. G. et al. Global patterns of 16S rRNA diversity at a depth of millions of sequences per sample. Proc Natl Acad Sci U S A 108 **Suppl 1**, 4516–4522, doi:10.1073/pnas.1000080107 (2011).

89. Bolyen, E. et al. Reproducible, interactive, scalable and extensible microbiome data science using QIIME 2. Nature Biotechnology 37, 852–857 (2019).

90. Callahan, B. J. et al. DADA2: High-resolution sample inference from Illumina amplicon data. Nature Methods 13, 581–583, doi:10.1038/nmeth.3869 (2016).

91. Bokulich, N. A. et al. Optimizing taxonomic classification of marker-gene amplicon sequences with QIIME 2’s q2-feature-classifier plugin. Microbiome 6, 90, doi:10.1186/s40168-018-0470-z (2018).

92. Maurice, C. F. & Turnbaugh, P. J. Quantifying and Identifying the Active and Damaged Subsets of Indigenous Microbial Communities Methods in Enzymology. Methods in Enzymology 531, 91–107, doi:doi: 10.1016/B978-0-12-407863-5.00005-8. (2013).

93. Bolger, A. M., Lohse, M. & Usadel, B. Trimmomatic: a flexible trimmer for Illumina sequence data. Bioinformatics 30, 2114–2120 (2014).

94. Bankevich, A. et al. SPAdes: a new genome assembly algorithm and its applications to single-cell sequencing. Journal of computational biology 19, 455–477, doi:10.1089/cmb.2012.0021 (2014).

95. Roux, S., Enault, F., Hurwitz, B. L. & Sullivan, M. B. VirSorter: mining viral signal from microbial genomic data. PeerJ 3, e985, doi:10.7717/peerj.985 (2015).

96. Fu, L., Niu, B., Zhu, Z., Wu, S. & Li, W. CD-HIT: accelerated for clustering the next- generation sequencing data. Bioinformatics 28, 3150–3152, doi:10.1093/bioinformatics/bts565 (2012).

97. Guerin, E. et al. Biology and Taxonomy of crAss-like Bacteriophages, the Most Abundant Virus in the Human Gut. Cell Host Microbe 24, 653–664.e656, doi:10.1016/j.chom.2018.10.002 (2018).

98. Langmead, B. & Salzberg, S. L. Fast gapped-read alignment with Bowtie 2. Nat Methods 9, 357–359, doi:10.1038/nmeth.1923 (2012).

99. Li, H. et al. The Sequence Alignment/Map format and SAMtools. Bioinformatics 25, 2078–2079, doi:10.1093/bioinformatics/btp352 (2009).

100. Roux, S., Emerson, J. B., Eloe-Fadrosh, E. A. & Sullivan, M. B. Benchmarking viromics: an in silico evaluation of metagenome-enabled estimates of viral community composition and diversity. PeerJ 5, e3817, doi:10.7717/peerj.3817 (2017).

