## additional file 1 for "Transplantation of bacteriophages from ulcerative colitis patients shifts the gut bacteriome and exacerbates severity of DSS-colitis"

**SUPPLEMENTARY FIGURES**

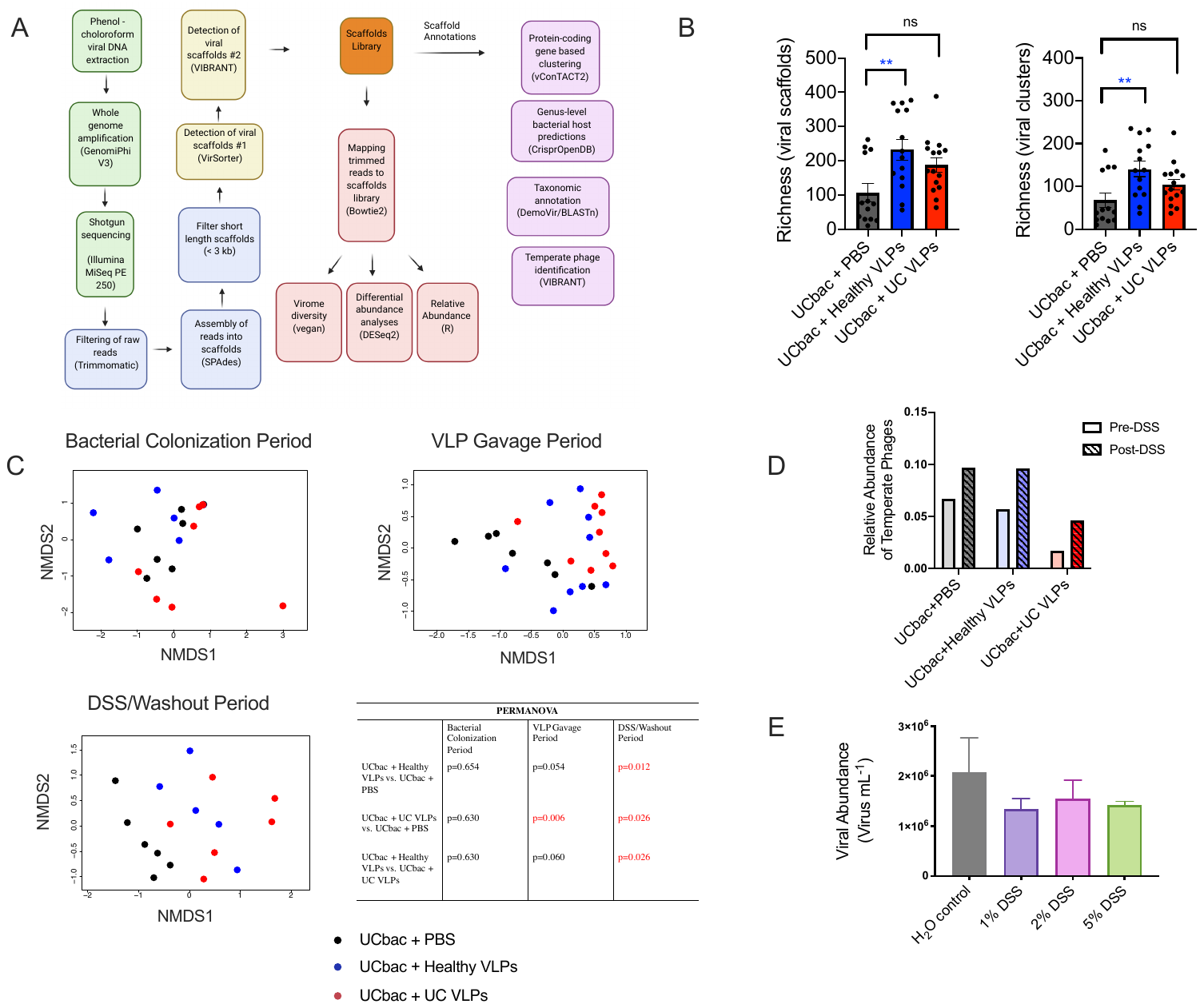

**Supplementary Figure S1.** **Virome analyses on stock and mouse VLPs**. **Related to Figure 2 and Figure 5. (**A) Viral metagenomic analyses was performed on fecal pellets from UC-HMA mice and human pooled UC or healthy VLP stocks. Viral scaffolds were detected using VirSorter and VIBRANT from mice and human samples to form a non-redundant scaffolds library. Data shown in (B-D) refers to experiment in Figure 1B (trial #1). Mouse fecal samples in each cage were pooled from 2 mice (n=3 cages per treatment group, 2 mice per cage). (B) Mean viral richness of scaffolds and VCs was determined between mice given healthy VLPs, UC VLPs or PBS. All samples after the first dose of VLPs or PBS given to HMA mice were included for analyses. Significance was assessed using one-way ANOVA using Tukey’s multiple comparison test (**p $\leq$ 0.01). (C) NMDS of Bray-Curtis dissimilarity of VCs between HMA mice given healthy VLPs, UC VLPs or PBS during the bacterial colonization period, VLP gavage period or the DSS washout period. (Bottom-right) adonis PERMANOVA table, assessing significant differences (p $\leq$ 0.05) in Bray-Curtis dissimilarity between HMA mice given healthy VLPs, UC VLPs or PBS. Samples from all time points of the longitudinal experiment were included in the NMDS and comparative analyses. Dots represent pooled mouse fecal samples at a single time point. UCbac, UC-HMA mice. (D) Relative abundance of scaffolds identified as temperate before and after DSS-administration. (E) Mean VLP abundance from the supernatant of bacterial cultures from pooled UC fecal samples grown *in vitro*. Bacteria were grown anaerobically in triplicate at 37°C in BHI media supplemented with hemin (5 μg/mL) and vitamin K (1 μg/mL). DSS or H_2_O were added to cultures at early exponential phase (0.25-3 OD_600_) and VLP supernatant was sampled for enumeration at stationary phase after 14 hr of growth.

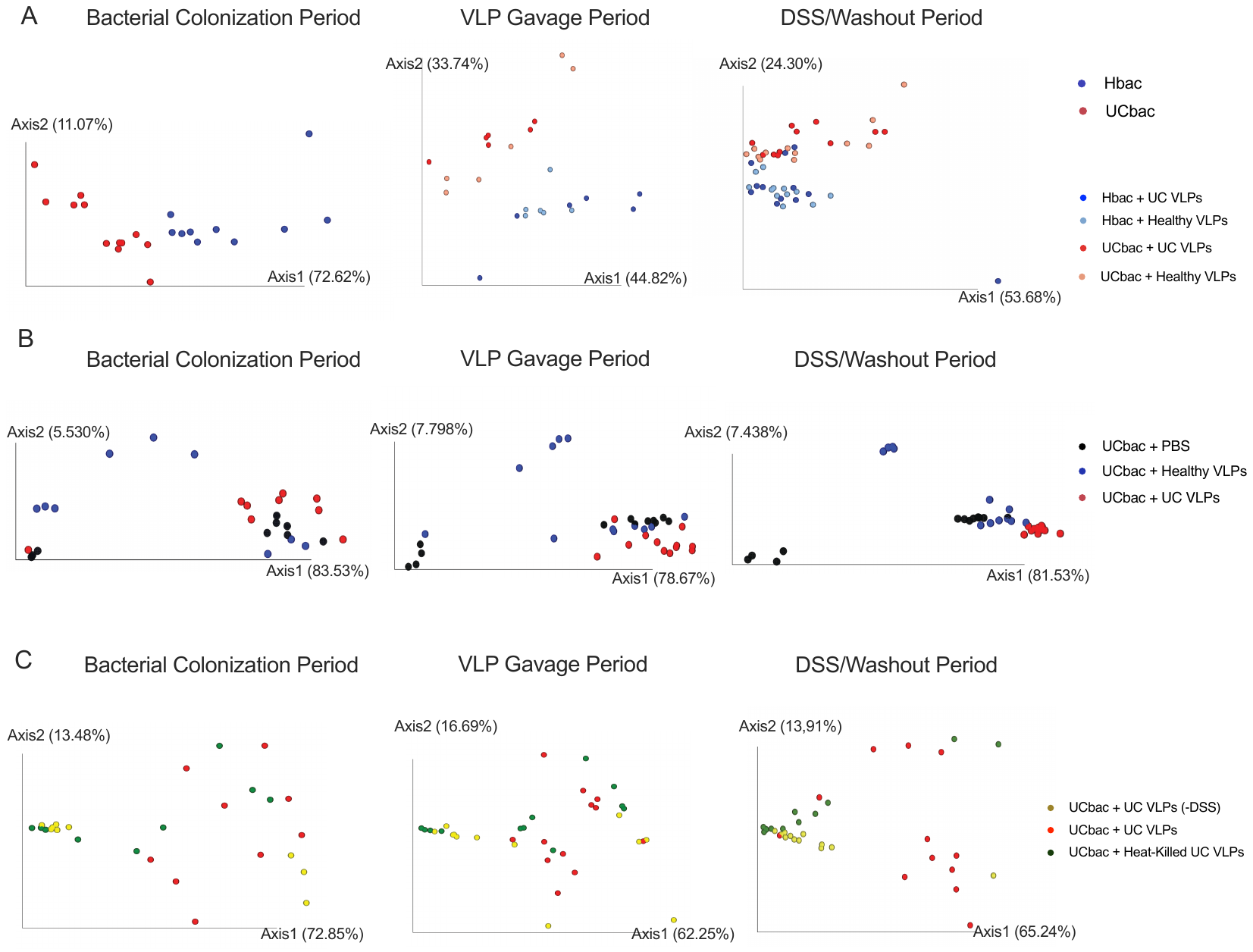

**Supplementary Figure S2. PcoA on Weighted UniFrac distances in HMA mice given VLPs**. **Related to Figure 3 and Figure 6.** (A) UC and Healthy-HMA mice were given a single dose of healthy or UC VLPs. Refers to experiment in Figure 1A. (B) UC-HMA mice were given 4 doses of healthy VLPs, UC VLPs, or PBS. Data shown is from the second of two independent trials. Data refers to experiment in Figure 1B (trial #2). **(**C) UC-HMA or GF mice were given 4 doses of UC VLPs or heat-killed UC VLPs. Data refers to experiment in Figure 1C. Mouse fecal samples in each cage were pooled from 1 or 2 mice (n=3 cages per treatment group, 1 or 2 mice per cage). Dots represent pooled mouse fecal samples at a single time point. Hbac, healthy-HMA mice; UCbac, UC-HMA mice. Samples from all time points of each experiment were included in the PcoA and comparative analyses.

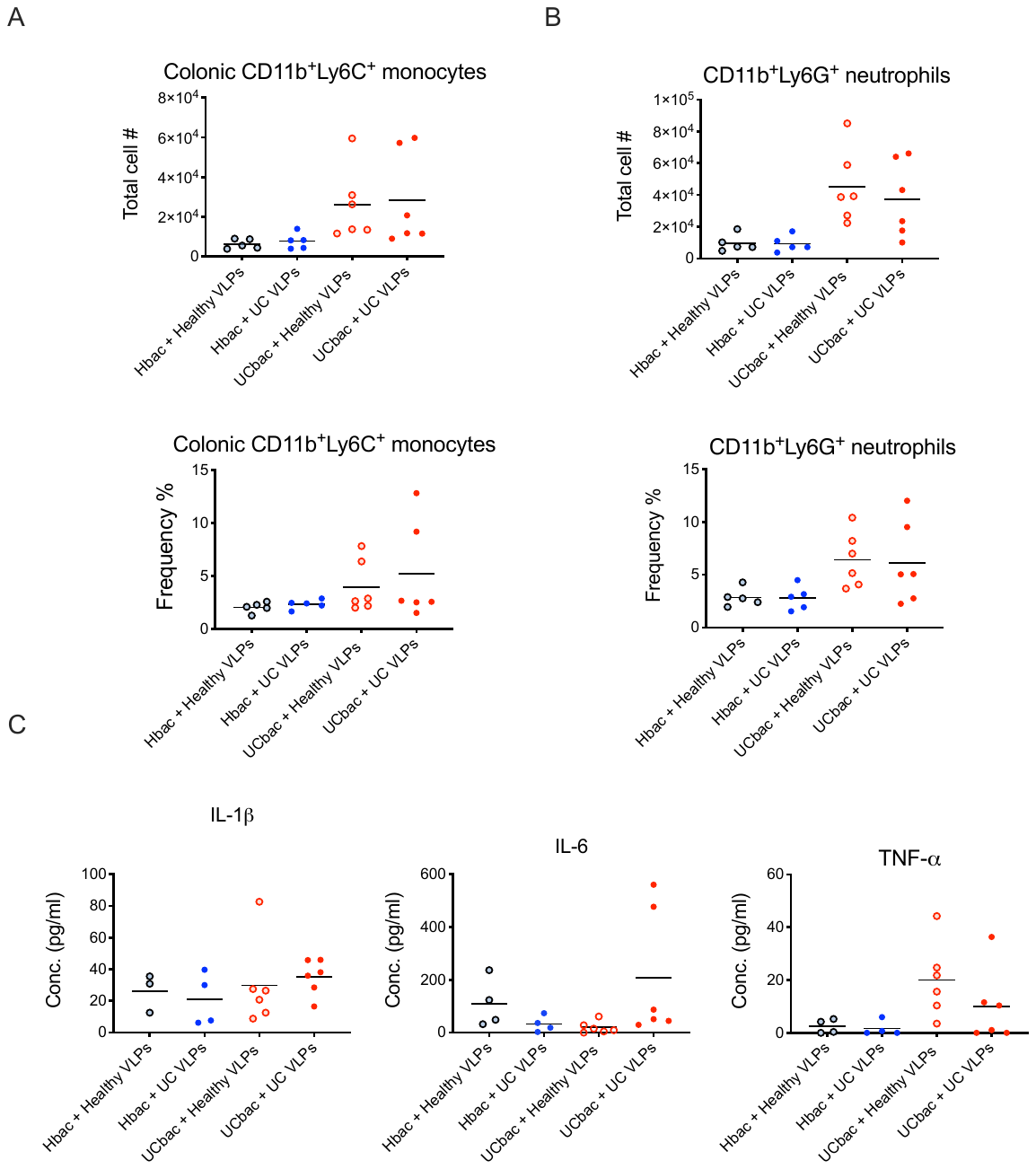

**Supplementary Figure S3. Experimental colitis severity of UC-HMA and healthy-HMA mice given a single dose of healthy or UC VLPs.** **Related to Figure 3.** (A) Mean absolute number and frequency of inflammatory monocytes (CD11b+Ly6C+Ly6G-) isolated from the colon at day 10 post-DSS administration. (B) Mean absolute number and frequency of neutrophils (CD11b+Ly6C-Ly6G+) isolated from the colon at day 10 post-DSS administration. (C) Mean Inflammatory cytokine production from colon tissue explants at day 10 post-DSS administration. Data were analyzed by one-way ANOVA, using Tukey’s multiple comparison test. Dots represent individual mice. Hbac + healthy/UC VLPs, n = 4 mice per group; UCbac + healthy/UC VLPs, n=6 mice per group. Hbac, healthy-HMA mice; UCbac, UC-HMA mice.

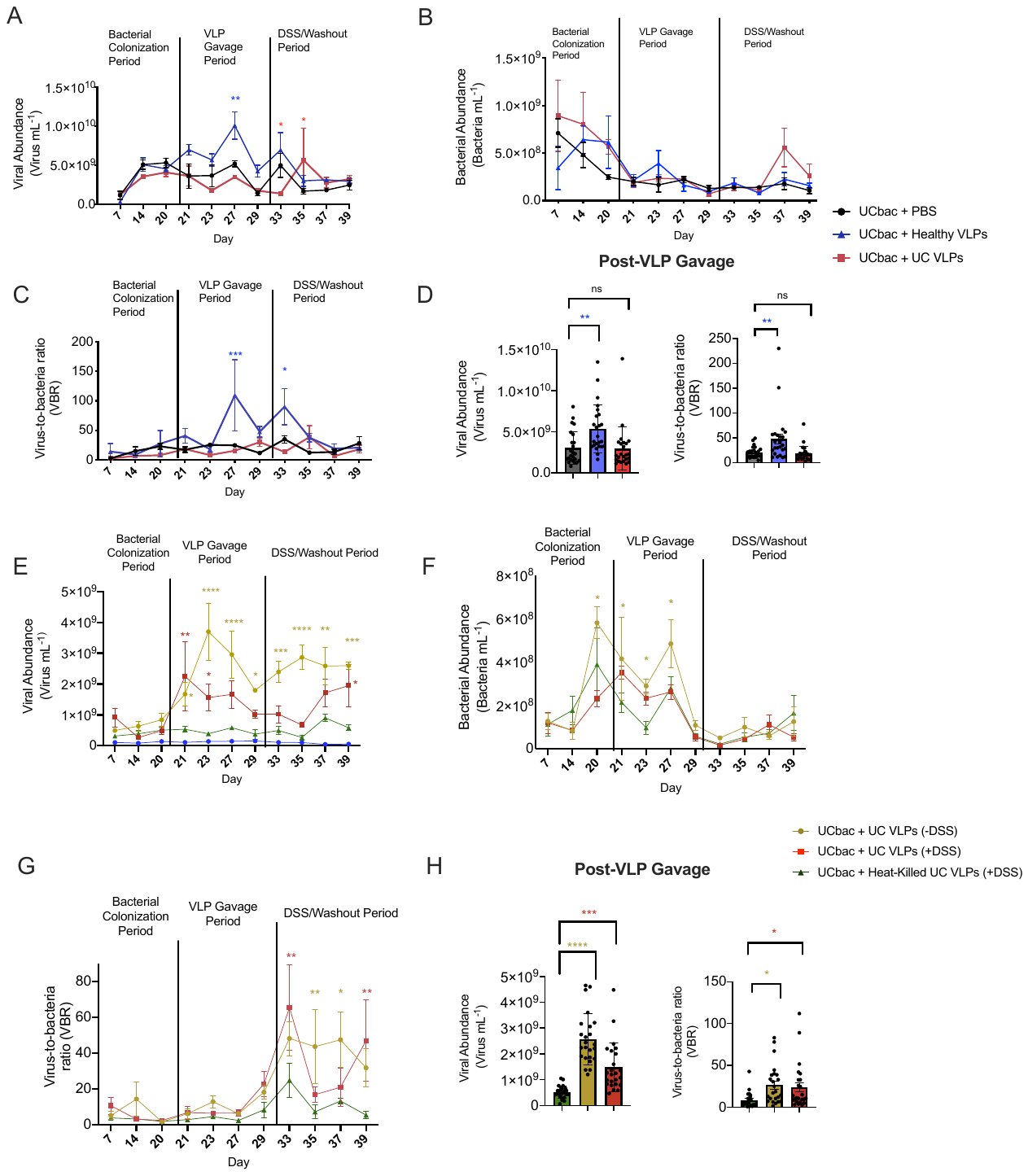

**Supplementary Figure S4. Viral and bacterial abundance in UC-HMA mice**. **Related to Figure 4.** Viral abundance was determined from mouse fecal pellets using epifluorescence microscopy and compared to bacterial abundances obtained by flow cytometry after staining with SybrGREEN I to obtain VBRs. Mean total viral abundance and mean total VBR post-VLP gavage were compared between treatment groups after the first dose of VLPs, heat-killed VLPs, or PBS was given to mice. (A-D) Red and blue asterisks indicate significant differences between the PBS control and HMA mice given UC VLPs or healthy VLPs respectively. (E-F) Gold and red asterisks indicate significant differences between heat-killed controls and HMA mice given UC VLPs (-DSS) or UC VLPs (+DSS) respectively. Data shown in A-D is from the second of two independent trials. Mouse fecal samples in each cage were pooled from 2 mice (n=3 cages per treatment group). Significance at each time point was assessed using Dunnett’s multiple comparisons test (*p $\leq$ 0.05, **p $\leq$ 0.01, ***p $\leq$ 0.001, ****p $\leq$ 0.0001). Significance post-VLP gavage was assessed using one-way ANOVA and Tukey’s multiple comparison test (*p $\leq$ 0.05, ***p $\leq$ 0.001, ****p $\leq$ 0.0001). Dots represent abundance or VBR from pooled mouse fecal samples at a single time point. Error bars, SE. UCbac, UC-HMA mice.

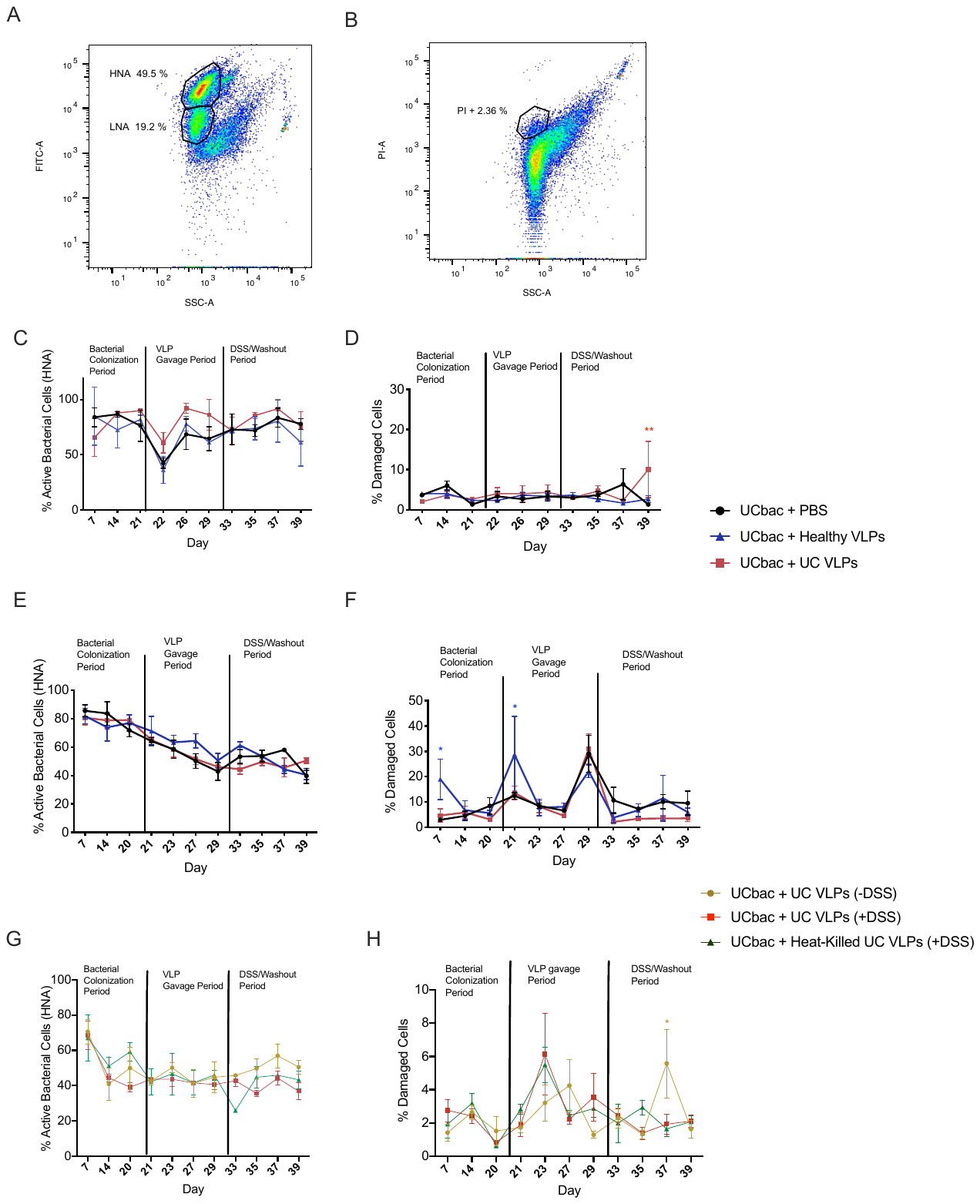

**Supplementary Figure S5. Bacterial physiology of HMA mice**. **Related to Figure 4.** Bacterial communities were extracted from mouse fecal pellets under anaerobic conditions. To determine the proportion of active and damaged bacterial cells, fecal bacterial communities were stained with SybrGreen and PI respectively. (A) Gating strategy for mouse fecal bacteria to determine high nucleic acid (HNA) bacterial cells and low nucleic acid (LNA) bacterial cells. (B) Gating strategy to determine damaged bacterial cells using propidium iodide (PI). (C-F) Red and blue asterisks indicate significant differences between the PBS control and HMA mice given UC VLPs or healthy VLPs respectively. (G-H) Gold and red asterisks indicate significant differences between heat-killed controls and HMA mice given UC VLPs (-DSS) and UC VLPs (+DSS) respectively. Significance at each time point was assessed using Dunnett’s multiple comparisons test (*p $\leq$ 0.05, **p $\leq$ 0.01). At each time point, mouse fecal samples in each cage were pooled from 2 mice (n=3 cages per group, 6 mice per group). Dots represent active or damaged bacterial cells from pooled mouse fecal samples at a single time point. Error bars, SE. UC bac, UC HMA-mice.

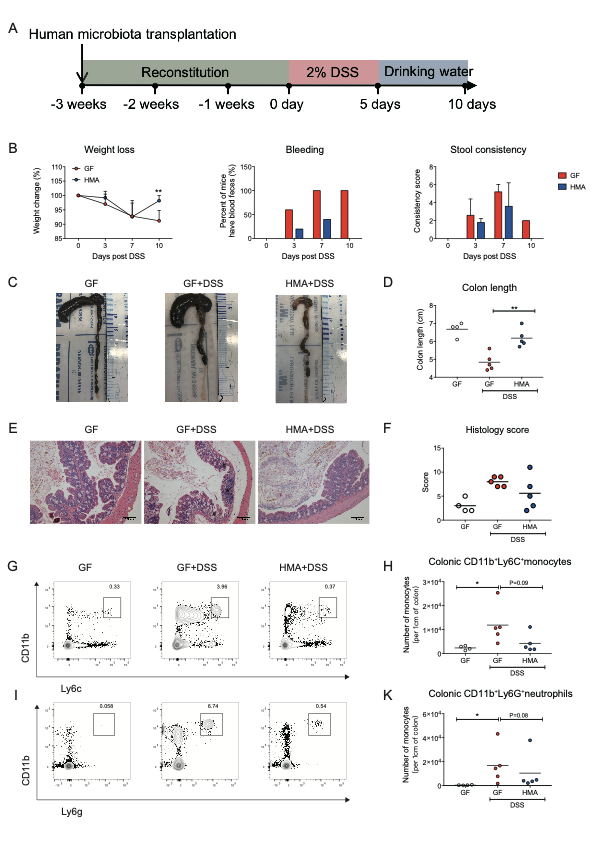

**Supplementary Figure S6. Human microbiota protects mice from experimental colitis.** **Related to Figure 7.** (A) Experimental design of DSS-colitis on HMA mice. (B) Changes of clinical disease activity index (DAI) including mean weight loss (left), mean GI tract bleeding (middle), and mean stool consistency (right) following DSS treatment. n=5 mice for each group. (C) Gross picture of colons at day 10 post DSS administration. (D) Mean length of colon of Control GF mice without DSS (white), GF mice (red) and HMA mice (blue) at day 10 post DSS administration. (E) Representative H&E staining of paraffin-embedded colon tissue at day 10 post DSS administration (scale bars are 100μm). Asterisk (*) indicates area of cellular infiltration; Number sign (#) indicates area of distortion of crypt architecture. (F) Mean histopathology of the effect of human gut microbiota colonization on DSS-induced colon damage in mice at day 10 post DSS administration. (G) Representative contour plots of colon monocytes defined as Viability-CD45+Ly6G-CD11b+Ly6C+ cells. (H) Mean total cell numbers of colon monocytes at day 10 post DSS administration. (I) Representative contour plots of colon neutrophils defined as Viability-CD45+CD11b+Ly6G+ cells. (K) Mean total cell numbers of colon neutrophils at day 10 post DSS administration. B-K. Data shown from one experiment. Each dot represents an individual mouse. Data were analyzed by two-way ANOVA with Bonferroni for multiple comparisons for panel B; Data were analyzed using an two-tailed unpaired parametric t test (*p < 0.05, **p < 0.01, ***p < 0.001) for panels D, H, K. Error bars represent the SD.

**SUPPLEMENTARY TABLES**

| **PERMANOVA** | | | |
| --- | --- | --- | --- |
| **Experiment referenced in Fig. 1A** | | | |
|  |  | **VLP Gavage period** | **DSS/Washout**  **Period** |
| Hbac + Healthy VLPs vs. Hbac + UC VLPs |  | p=0.307 | p=0.8420 |
| UCbac + Healthy VLPs vs. UCbac + UC VLPs |  | p=0.577 | p=0.8420 |
| **Experiment referenced in Fig. 1B, trial #2** | | | |
|  | **Bacterial Colonization Period** | **VLP**  **Gavage**  **Period** | **DSS/Washout Period** |
| UCbac + Healthy VLPs vs. UCbac + PBS | p=0.398 | p=0.376 | p=0.063 |
| UCbac + Healthy VLPs vs. UCbac + UC VLPs | p=0.246 | p=0.001 | p=0.0015 |
| UCbac + PBS vs. UCbac + UC VLPs | p=0.171 | p=0.003 | p=0.0015 |
| **Experiment referenced in Fig. 1C** | | | |
|  | **Bacterial Colonization Period** | **VLP Gavage period** | **DSS/Washout Period** |
| UCbac+ heat-killed UC VLPs vs. UCbac+ UC VLPs | p=0.069 | p=0.129 | p=0.003 |
| UCbac+ heat-killed UC VLPs vs. UCbac+ UC VLPs (- DSS) | p=0.500 | p=0.252 | p=0.161 |
| UCbac+ UC VLPs vs. UCbac + UC VLPs (- DSS) | p=0.090 | p=0.129 | p=0.003 |

**Supplementary Table. S2 PERMANOVA on weighted UniFrac distances between UC or healthy-HMA mice given healthy VLPs or UC VLPs. Related to Figure 3 and Figure 6.** Mouse fecal samples in each cage were pooled from 1 or 2 mice (n=3 cages per treatment group, 1 or 2 mice per cage). Hbac, healthy-HMA mice; UCbac, UC-HMA mice.

| **Species** | **Pairwise Difference** |
| --- | --- |
| *Anaerotruncus* sp. (2) trial #1 | Decreased in UC VLP treatment |
| *Ruminiclostridium 5* sp. (1) trial #1 | Decreased in UC VLP treatment |
| *Sellimonas* sp. (3) trial #1 | Increased in UC VLP treatment |
| *Eubacterium limosum* (2) trial #1 | Decreased in UC VLP treatment |
| *Clostridium sensu stricto* 1 sp. (1) trial #2 | Increased in PBS control treatment |
| *Enterococcus* sp. (1) trial #2 | Decreased in PBS control treatment |
| [*Eubacterium*] *fissicatena* group sp. (3) trial #2 | Increased in healthy VLP treatment |
| *Negativibacillus* sp. (3) trial #2 | Increased in UC VLP treatment |
| Ruminococcaceae UCG-005 sp. (3) trial #2 | Increased in UC VLP treatment |
| *Epulopiscium* sp. (1) trial #2 | Decreased in UC VLP treatment |
| [*Eubacterium*] *coprostanoligenes* sp. (2) trial #2 | Increased in PBS control treatment |

**Supplementary Table. S5 Differentially abundant species during VLP gavage period in HMA mice given healthy VLPs, UC VLPs, or PBS.** **Related to Figure 6.**

Taxonomy was assigned using the Qiime2 feature classifier and differentially abundant species were determined using analysis of the composition of microbes (ANCOM). Differentially abundant species were identified during the VLP gavage period. Taxa that were found to be differentially abundant during bacterial colonization were not included. Numbers in parentheses (1-3) correspond to likelihood that pairwise differences were due to phage treatment or isolator effect (see STAR methods for ranking criteria). Data shown is aggregated from two independent trials. In each experiment, mouse fecal samples in each cage were pooled from 2 mice (n=3 cages per treatment group).

| **Species** | **Pairwise Difference** |
| --- | --- |
| *Sellimonas* sp. (3) trial #1 | Increased in UC VLP treatment |
| *Eubacterium limosum* (2) trial #1 | Decreased in UC VLP treatment |
| *Ruminiclostridium 5* sp. (1) trial #1 | Decreased in UC VLP treatment |
| *Anaerotruncus* sp. (2) trial #1 | Decreased in UC VLP treatment |
| *Enterococcus* sp. (1) trial #2 | Decreased in PBS control treatment |
| [*Eubacterium*] *coprostanoligenes* group sp. (2) trial #2 | Increased in PBS control treatment |
| [*Eubacterium*] *fissicatena* group sp. (3) trial #2 | Increased in healthy VLP treatment |
| *Butyricicoccus* sp.(2) trial #2 | Increased in healthy VLP treatment |
| *Negativibacillus* sp. (3) trial #2 | Increased in UC VLP treatment |
| *Parabacteroides distasonis* (2) trial #2 | Increased in healthy VLP treatment |
| *Ruminococcaceae* UCG-005 sp. (3) trial #2 | Increased in UC VLP treatment |
| *Clostridium sensu stricto* 1 sp. (1) trial #2 | Increased in PBS control treatment |
| Uncultured *Clostridium* sp. (2) trial #2 | Increased in healthy VLP treatment |
| *Anaerotruncus* sp. (3) trial #2 | Increased in UC VLP treatment |
| *Tyzzerella* sp. (3) trial #2 | Increased in UC VLP treatment |
| *Candidatus Stoquefichus* (2) trial #2 | Increased in healthy VLP treatment |
| *Flavonifractor* sp. (3) trial #2 | Increased in UC VLP treatment |
| *coprostanoligenes* UBA1819 sp. trial #2 (3) | Increased in UC VLP treatment |
| *Eubacterium dolichum* trial #2 (3) | Decreased in UC VLP treatment |

**Supplementary Table. S6 Differentially abundant species during DSS/washout period in HMA mice given healthy VLPs, UC VLPs, or PBS**. **Related to Figure 6.** Taxonomy was assigned using the Qiime2 feature classifier and differentially abundant species between treatment groups were determined using analysis of the composition of microbes (ANCOM). Differentially abundant species were identified during the DSS/washout period. Taxa that were found to be differentially abundant during bacterial colonization were not included. Numbers in parentheses (1-3) correspond to likelihood that pairwise differences were due to VLP treatment or isolator effect (see STAR methods for ranking criteria). Data shown is aggregated from two independent trials. In each experiment, mouse fecal samples in each cage were pooled from 2 mice (n=3 cages per treatment group).

| **Species** | **Pairwise Difference** |
| --- | --- |
| *Blautia hydrogenotrophica* (3) | Increased in heat-killed UC VLP treatment |
| *Escherichia*–*Shigella* sp. (1) | Decreased in heat-killed UC VLP |
| [*Eubacterium*] *fissicatena* group sp. (3) | Increased in heat-killed UC VLP treatment |

**Supplementary Table. S7 Differentially abundant species during VLP gavage period in HMA mice given UC VLPs (+/- DSS), or heat-killed UC VLPs. Related to Figure 6.** Taxonomy was assigned using the Qiime2 feature classifier and differentially abundant species between treatment groups were determined using analysis of the composition of microbes (ANCOM). Differentially abundant species were identified during the VLP gavage period. Taxa that were found to be differentially abundant during the bacterial colonization period were not included. Numbers in parentheses (1-3) correspond to likelihood that pairwise differences were due to VLP treatment or isolator effect (see STAR methods for ranking criteria). Mouse fecal samples in each cage were pooled from 2 mice (n=3 cages per treatment group).

| **Species** | **Pairwise Difference** |
| --- | --- |
| *Blautia hydrogenotrophica* (3) | Increased in heat-killed UC VLP treatment |
| *Lachnospiraceae* sp. (1) | Decreased in heat-killed UC VLP treatment |
| *Alistipes* sp. (2) | Decreased in heat-killed UC VLP treatment |
| *Escherichia*–*Shigella* sp. (1) | Decreased in heat-killed UC VLP treatment |
| [*Eubacterium*] *fissicatena* group sp. (3) | Increased in heat-killed UC VLP treatment |
| Uncultured *Clostridium* sp. (2) | Increased in heat-killed UC VLP treatment |

**Supplementary Table. S8 Differentially abundant species during DSS/ washout period in HMA mice given UC VLPs (+/- DSS) or heat-killed UC VLPs. Related to Figure 6.** Taxonomy was assigned using the Qiime2 feature classifier and differentially abundant species between treatment groups were determined using analysis of the composition of microbes (ANCOM). Taxa that were found to be differentially abundant during the bacterial colonization period were not included. Numbers in parentheses (1-3) correspond to likelihood that pairwise differences were due to VLP treatment or isolator effect (see STAR methods for ranking criteria). Mouse fecal samples in each cage were pooled from 2 mice (n=3 cages per treatment group).
